# *Corynebacterium diphtheriae* causes keratinocyte-intrinsic ribotoxic stress and NLRP1 inflammasome activation in a model of cutaneous diphtheria

**DOI:** 10.1101/2023.01.16.524188

**Authors:** Kim S. ROBINSON, Gee Ann TOH, Muhammad Jasrie FIRDAUS, Khek Chian THAM, Pritisha ROZARIO, Chrissie LIM, Ying Xiu TOH, Zhi Heng LAU, Sophie Charlotte BINDER, Jacob MAYER, Carine BONNARD, Florian I. SCHMIDT, John E. A. COMMON, Franklin L. ZHONG

## Abstract

NLRP1 is an innate immune sensor protein that activates inflammasome-driven pyroptotic cell death. Recent work demonstrates that human NLRP1 has evolved to sense viral infections. Whether and how human NLRP1 responds to other infectious agents is unclear. Here, and in a companion manuscript, we report that human NLRP1, as an integral component of the ribotoxic stress response (RSR), is activated by bacterial exotoxins that target human ribosome elongation factors EEF1 and EEF2, including Diphtheria Toxin (DT) from *Corynebacterium diphtheria*e, exotoxin A from *Pseudomonas aeruginosa* and sidI from *Legionella pneumophila*. In human keratinocytes, DT activates RSR kinases ZAKα, p38 and JNKs, upregulates a set of signature RSR transcripts and triggers rapid NLRP1-dependent pyroptosis. Mechanistically, these processes require 1) DtxR-mediated de-repression of DT production in the bacteria, as well as 2) diphthamide synthesis and 3) ZAKα/p38-driven NLRP1 phosphorylation in the host. In 3D human skin cultures, *Corynebacterium diphtheria*e infection disrupts barrier function and induces IL-1 driven inflammation. Pharmacologic inhibition of p38 and ZAKα suppresses keratinocyte pyroptosis and rescues barrier integrity of *Corynebacterium diphtheria*e-treated organotypic skin. In summary, these findings implicate RSR and the NLRP1 inflammasome in antibacterial innate immunity and might explain certain aspects of diphtheria pathogenesis.

**KEY POINTS:**

1. EEF1/EEF2-targeting bacterial exotoxins activate the human NLRP1 inflammasome.
2. DT+ve toxigenic *Corynebacterium diphtheriae* induces ZAKα-driven RSR and NLRP1-driven pyroptosis in human keratinocytes.
3. Identification of transcripts that are induced by multiple RSR agents across multiple cell types.
4. p38 and ZAKα inhibition rescues epidermal integrity by limiting pyroptosis in 3D skin mode of cutaneous diphtheria.

## INTRODUCTION

Before the introduction of a worldwide vaccination campaign, diphtheria, caused by the Gram positive extracellular bacterium *Corynebacterium diphtheria*e, remained a deadly scourge throughout modern human history, with a case fatality rate up to 10% in adults and 20% in children under five (Sharma et al., 2019; Acosta and Tiwari, 2020). The respiratory form of diphtheria causes tissue necrosis of tonsils, throat, and larynx, leading to the formation of ‘pseudomembrane’ - an eponymous hallmark symptom (latin, *diphthera*, ‘leather hide’). Fatality is typically associated with respiratory failure as well as myocarditis, polyneuropathy and kidney damage caused by systemic dissemination of bacterial toxins. *C. diphtheria*e, and related C. *ulcerans* also infect skin wounds and cause cutaneous diphtheria, which is characterized by deep ulcers and extensive epidermal and dermal necrosis (Hadfield et al., 2000).

As early as 1888, Roux and Yersin and others discovered that the sterile filtrate of *C. diphtheria*e culture broth was sufficient in eliciting most of the diphtheria symptoms when injected into small animals. This was the first demonstration that diphtheria, in the strictest sense, is a toxin-mediated disease. Subsequent work demonstrated that a single secreted toxin, named diphtheria toxin (DT) was responsible for diphtheria pathogenesis (Holmes, 2000). These landmark discoveries paved the way for modern day diphtheria treatment and prevention. When administered early, antisera against DT together with antibiotics are a curative treatment for diphtheria. Vaccines that elicit a humoral immune response against DT, rather than the bacterium itself, have proven highly effective and are a cornerstone component of childhood vaccinations across the world (reviewed in (Murphy). As a result, worldwide incidence of diphtheria has been low in the last few decades, but localized outbreaks continue to occur, especially in overcrowded settings with inadequate vaccine coverage (Blumberg et al., 2018; Wise, 2022).

DT is a typical A/B exotoxin: DTB binds to the host receptor human pro-HBEGF and delivers DTA into the cytosol, where DTA catalyzes the conjugation of an ADP-ribose moiety to a post-translationally acquired unusual amino acid known as diphthamide at position of 715 of human Eukaryotic Translation Elongation Factor 2 (EEF2) (Holmes, 2000). This reaction inactivates EEF2 and shuts down host protein synthesis. Due to its potent toxicity, DT has now been successfully developed as a toxin payload in antibody conjugates for cancer treatment (Frankel et al., 2002). DT is also widely used as a research tool to carry out cell ablation experiments in engineered mouse models (Palmiter et al., 1987). As such, most of the biological effects of DT have been studied in the context of apoptosis, and it is currently unclear whether the human innate immune system directly responds to DT.

Here, and in a companion manuscript, we demonstrate that DT and other EEF1/EEF2-targeting bacterial toxins cause pyroptosis of primary human epithelial cells. Using primary keratinocyte and 3D reconstructed skin as a model of cutaneous diphtheria, we found that this process occurs by cell intrinsic activation of ZAKα-driven ribotoxic stress and the inflammasome sensor NLRP1. Pharmacologic inhibitors of ZAKα and p38 kinases prevent epidermal damage by purified and toxigenic *C. diphtheria*. Our results define DT as a bona fide ribotoxic stress inducer and a species-specific trigger for IL-1 driven inflammation in human epithelia. These results also establish a role of RSR and the human NLRP1 inflammasome in the pathogenesis of diphtheria and likely other bacterial infections.

## RESULTS

### EEF1/2-targeting bacterial toxins, DT, exotoxin A and sidI activate the ribotoxic stress and induce pyroptosis

NLRP1 is a versatile sensor for the cytosolic inflammasome complex(Martinon et al., 2002; Mitchell et al., 2019; Bachovchin, 2021). The human NLRP1 inflammasome was recently found to sense ribotoxic stress inducing agents which cause ribosome stalling and/or collisions, e.g. UVB and anisomycin (Robinson et al., 2022; Jenster et al., 2023). This discovery prompted us to investigate whether bacterial virulence factors that are known to affect host translation could similarly activate human NLRP1. We tested three secreted bacterial exotoxins, *C. diphtheriae* DT, *P. aeruginosa* exotoxin A (exoA) and *L. pneumophila* sidI (Fig. 1A). DT and exoA are ADP-ribosylating enzymes that inactivate EEF2 at a conserved diphthamide residue, whereas sidI acts as a tRNA mimic that inactivates EEF1 and ribosome components(Subramanian et al., 2022; Joseph et al., 2019). A small molecule inhibitor of EEF1, didemnin B (DDB, also known as Aplidine) (Crews et al., 1994) (Fig. 1A) and anisomycin were included as positive controls. SidI and DDB have recently been linked to the ribotoxic stress response (RSR) pathway downstream of ZAKα, but their effects on pyroptosis have not been shown(Subramanian et al., 2022).

**Figure 1.**
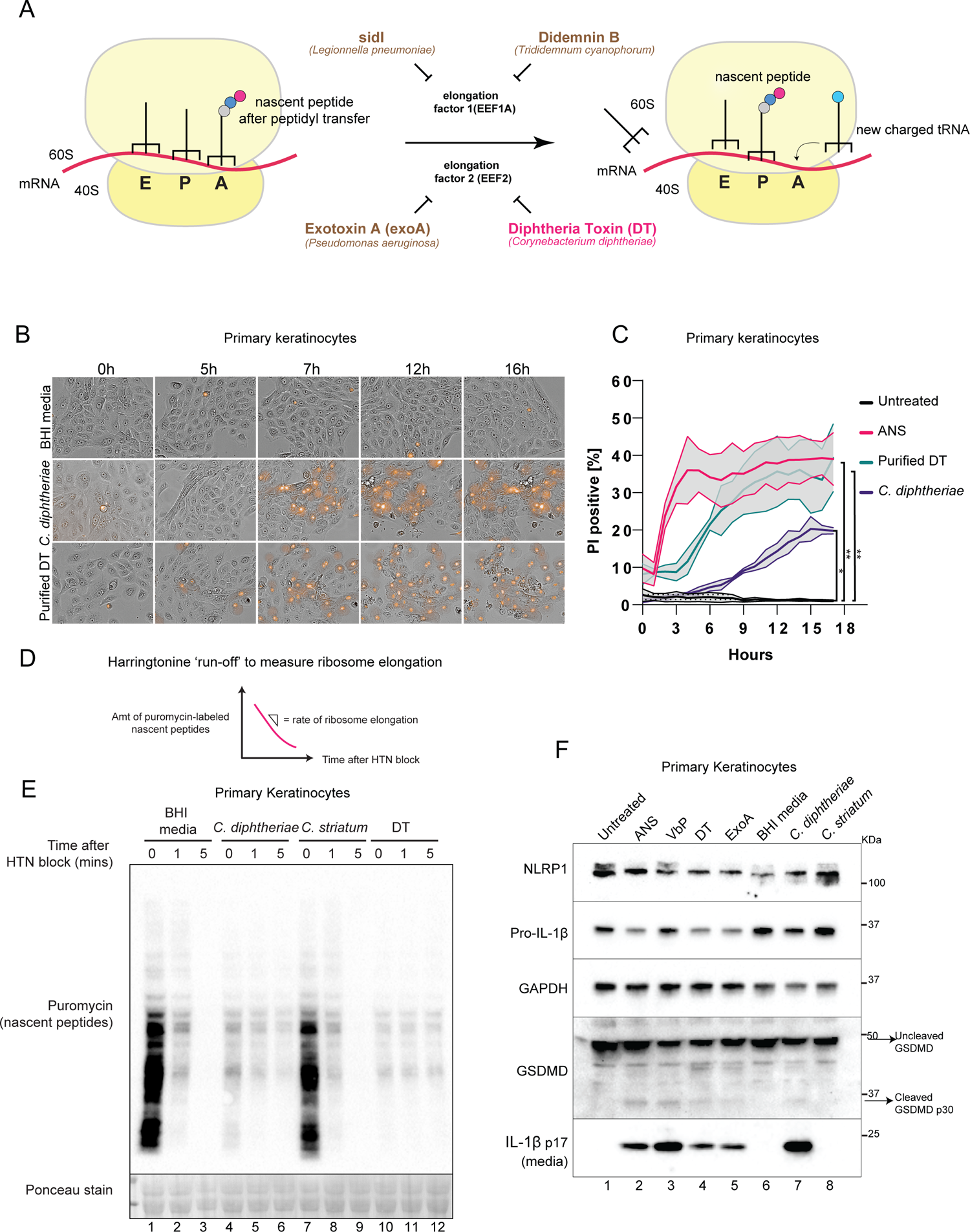
*C. diphtheriae*, DT and exoA activate pyroptosis in primary keratinocytes. (A) Simplified cartoon representation of eukaryotic ribosome translocation and EEF1/EEF2-targeting toxins. (B) Live cell imaging of primary keratinocytes treated with BHI media, sterile filtrates of *C. diphtheriae* grown in iron-restricted media or recombinant DT. Bright field images were overlaid with PI fluorescence (red). Images were taken at 20x magnification. (C) Quantification of the percentage of PI+ve cells over time. Error bars represent three biological replicates, with each drug treatment considered as one replicate. Significance values were calculated from Student’s t test at the 7 hour time point. (D) Principles of Harringtonine ribosome ‘run-off’ assay. (E) Anti-puromycin immunoblot of primary keratinocytes subjected to the Harringtonine run-off assay. Cells were treated with BHI media, sterile filtrates of *C. diphtheriae*, *C. striatum* for 6 hours or purified DT (150 ng/mL) for 3 hours before harringtonine addition (2µg/ml). Nascent peptides were then labeled with 10 µg/ml puromycin for 10 mins at different intervals. (F) Immunoblot of inflammasome activation markers in primary keratinocytes 24 hours post treatment. ANS, anisomycin (1 µM); VbP, Val-boro-Pro (3 µM); DT (150 ng/mL), exoA (1 µg/mL).

In primary keratinocytes, DT causes inflammasome-driven pyroptosis as shown by PI uptake (Fig. 1B, lower panel, and 1C), IL-1β p17 secretion and the processing of full length GSDMD into the p30 form (Fig. 1F). In immortalized N/TERT-1 (N/TERT) cells, DT also triggered inflammasome activation and pyroptosis, albeit with delayed kinetics and less potency (Fig. S1A). Exotoxin A induced minimal pyroptosis in N/TERT cells as measured by IL-1β secretion and GSDMD cleavage. These observations were likely due to the reduced expression of the DT and exoA entry receptors pro-HBEGF and LRP1 in N/TERT cells. TNFɑ significantly sensitized N/TERT cells both to DT and exoA-induced pyroptosis, in agreement with published findings that it upregulates LRP1 and pro-HBEGF (Morimoto et al., 1991; Morimoto and Bonavida, 1992). TNFɑ additionally increased the expression of IL-1B and NLRP1 itself; thus acting as a priming signal in N/TERT cells (Fig. S1A). Furthermore, bypassing the requirement of pro-HBEGF using an engineered anthrax factor conjugated DTA (LFn-DTA) also robustly induced pyroptosis in N/TERT cells (Milne et al., 1995) (Fig. S1B and S1C). Cell permeable EEF1 inhibitor DDB also triggered rapid pyroptosis in N/TERT cells in the absence of any priming signal (Fig. S2A and S2B). Transient overexpression of sidI, but not its glycosylase-defective mutant induced the formation of ASC-GFP specks in 293T-ASC-GFP-NLRP1 reporter cell line (Fig. S2C and S2D). Together, these data demonstrate that multiple EEF1 and EEF2 toxins can activate the human NLRP1 inflammasome.

### *Corynebacterium diphtheriae* induces ZAKα-driven RSR and NLRP1 inflammasome activation in human keratinocytes

Next we tested the effect of DT+ve pathogenic *C. diphtheriae* on primary keratinocytes. In all clinical *C. diphtheriae* strains, DT is encoded on a prophage under the control of an iron responsive repressor known as DtxR (Holmes, 2000). In agreement with previous findings, the secretion of DT from the toxigenic *C. diphtheriae* strain was detectable, but low, in iron-containing basal media (Fig. S3A-B). Iron chelation by 2,2′-dipyridyl dramatically induced the secretion of DT within 4 hours (Fig. S3A and S3C) in toxigenic *C. diphtheriae*, but not non-toxigenic (i.e. DT-ve) *C. diphtheriae* or related DT-ve *Corynebacterium* species. Using an Harringtonine ribosome ‘run-off’ assay (Schmidt et al., 2009) (Fig. 1D), we observed that similar to recombinant DT, the conditioned media of iron-restricted *C. diphtheriae* caused significant inhibition of ribosome elongation rates in primary human keratinocytes, an expected consequence of EEF2 inactivation (Fig. 1E). This was followed by pyroptotic cell death, as evident in IL-1β secretion, GSDMD cleavage and PI uptake (Fig. 1B, middle panel, Fig. 1F). The ability of *C. diphtheriae* to cause pyroptosis was dependent on iron depletion, as *C. diphtheriae* grown in iron-containing media was significantly attenuated at inducing IL-1β secretion and GSDMD cleavage (Fig. S3D). A related, nontoxigenic *Corynebacterium* species, *C. striatum,* failed to elicit pyroptosis in primary keratinocytes (Fig. 1F, lane 8). These results are consistent with historical findings by Yersin and Roux that sterile filtrates of toxigenic *C. diphtheriae* are cytotoxic in vitro and sufficient to cause diphtheria-like symptoms in susceptible, non-rodent mammals (reviewed in (Murphy). Our results provide a potential mechanism for these earlier observations and suggest that the main cytotoxic effect of *C. diphtheriae* on primary human epithelial cells is pyroptosis. At first glance, this may seem at odds with the often cited role of DT as an inducer of apoptosis; however, it is important to note that most in vitro studies on DT in the past used cancer cell lines, which do not express a functional NLRP1 inflammasome.

To ascertain whether *C. diphtheriae*-driven pyroptosis is strictly downstream of EEF2 inactivation and ribosome stalling, we knocked out DPH1, an enzyme that is required to convert EEF2 His715 into diphthamide, in N/TERT cells (Liu et al., 2004) (Fig. S4A). DPH1 deficient cells were completely unable to secrete IL-1β (Fig. S4B) or undergo pyroptosis (Fig. 2A) in response to either purified DT or *C. diphtheriae*. Thus, the inactivation of EEF2 by DT is an essential upstream step for *C. diphtheriae*-induced pyroptosis in human keratinocytes.

**Figure 2.**
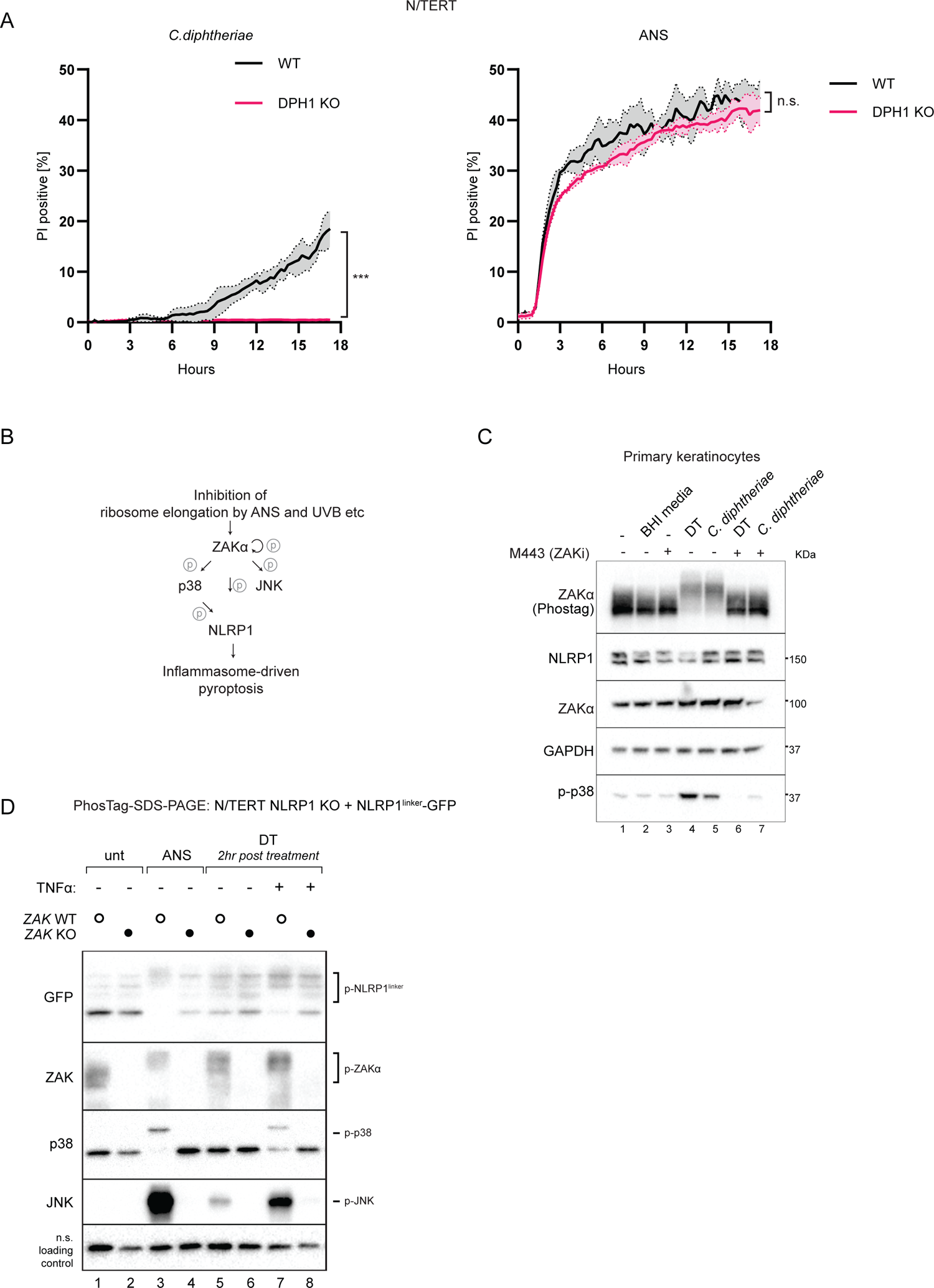
DT and *C. diphtheriae* trigger RSR downstream of EEF2 inactivation. (A) Kinetics of PI uptake in WT and DPH1 KO N/TERT cells treated with *C. diphtheriae* filtrate or ANS (1 µM) over 17 hours. Cells were primed with 25 ng/mL TNFɑ for 8 hours. Error bars represent three biological replicates, with each drug treatment considered as one replicate. Significance values were calculated from Student’s t test at the 7 hour time point. (B) Overview of ZAKɑ driven RSR pathway and RSR-dependent NLRP1 activation. (C) Immunoblot of ZAKɑ, p38 NLRP1 in primary keratinocytes treated with the indicated triggers in the presence of DMSO or M443 (1 µM). M443 were added 10 mins before DT or bacterial fitrate. (D) Immunoblot of RSR kinases and GFP-NLRP1^linker^ in control and ZAKɑ KO N/TERT cells.

Recently, MAP3K ZAKα was identified to be the long sought after RSR sensor that activates p38 and JNK in response to ribosome stalling and/or collisions (Vind et al., 2020b; Wu et al., 2020; Vind et al., 2020a). We further demonstrated that ZAKα, together with its downstream kinase p38, directly phosphorylates a species-specific linker domain on human NLRP1, a critical step in anisomycin and UVB-induced NLRP1 activation (Robinson et al., 2022; Jenster et al., 2023) (Fig. 2B). Even though the ability of DT to inhibit ribosome elongation is well established, it has not been studied as a direct inducer of the ZAKα-activated RSR. In primary keratinocytes treated with *C. diphtheriae* conditioned media or purified DT, we observed striking p38 phosphorylation and ZAKα auto-phosphorylation at 3 hours (Fig. 2C). In addition, in TNFα-primed N/TERT cells stably expressing a GFP-tagged NLRP1 linker, DT led to rapid NLRP1 linker phosphorylation (Fig. 2D). Genetic deletion of *ZAKα* or a small molecule inhibitor of ZAKα kinase activity, M443 abrogated all hallmarks of ribotoxic stress signaling, including the phosphorylation of ZAKα itself, p38, JNK and the NLRP1 linker domain (Fig. 2D). These results establish DT as a bona fide ribotoxic stress inducer, and suggest that *C. diphtheriae* likely activates the NLRP1 inflammasome downstream of ZAKα-driven ribotoxic stress induction. To test this genetically, we compared the response of control, NLRP1 KO and ZAKα KO primary human keratinocytes to recombinant DT, *C. diphtheriae* conditioned media, and known NLRP1 activators anisomycin (ANS) and Val-boro-Pro (VbP). CRISPR/Cas9-mediated ZAKα knockout abrogated PI uptake induced by *C. diphtheriae* and DT throughout the 16 hour time course (Fig. 3A and 3B, Fig. S5B and S5C), suggesting that the ZAKα-driven RSR is required for both fast-onset pyroptosis and late onset necrosis following DT intoxication. NLRP1 KO similarly abrogated PI uptake at early time points but did not affect late onset necrosis (Fig. 3C-D). ZAKα KO and NLRP1 KO cells were both unable to secrete mature IL-1β or cleave GSDMD into the p30 form after DT and *C. diphtheriae* treatment, as compared to Cas9 control cells (Fig. 3E, 3F, Fig. S5D). The ZAKα kinase inhibitor, M443 also blocked pyroptosis and inflammasome activation by DT (Markowitz et al., 2016), although its effect on late stage necrosis was less pronounced than the CRISPR/Cas9-mediated deletion of ZAKα (Fig. S5E, S6B). These data demonstrate that NLRP1, downstream of ZAKα, is indeed the main inflammasome sensor activated by *C. diphtheriae* and DT. Next, we examined the requirement of the p38 kinase using a clinical grade inhibitor Neflamapimod as well as p38 α/β double knockout (p38 DKO) N/TERT cells. SImilar to conventional ribotoxic stress inducers (ANS and UVB), p38 ɑ/β deletion or inhibition reduced DT-driven pyroptosis and IL-1β secretion in primed N/TERT cells, albeit to a lesser extent than ZAKα inhibition or deletion (Fig. S6A-C). These data also suggest that DT, similar to ANS, is unlikely to engage in the DPP9-dependent pathway of NLRP1 inflammasome activation (Robinson et al., 2022; Jenster et al., 2023). For additional mechanistic proof, we mutated one of the key ZAKα and p38 phosphorylation sites on the NLRP1 linker (Fig. 3G). We have previously shown that the resultant 3A mutant (T178A, S179A, T180A) is unable to execute ANS- and UVB-triggered pyroptosis, but does not affect VbP-induced NLRP1 activation (Robinson et al., 2022). Using NLRP1 KO N/TERT cells reconstituted with either wild-type or the NLRP1 3A mutant, we found that this site is also indispensable for DT- and exoA-driven pyroptosis (Fig. 3H, Fig. S6D and S6E). Collectively, these data show that DT activates the NLRP1 inflammasome via ZAKα-and p38-mediated phosphorylation of the NLRP1 linker.

**Figure 3.**
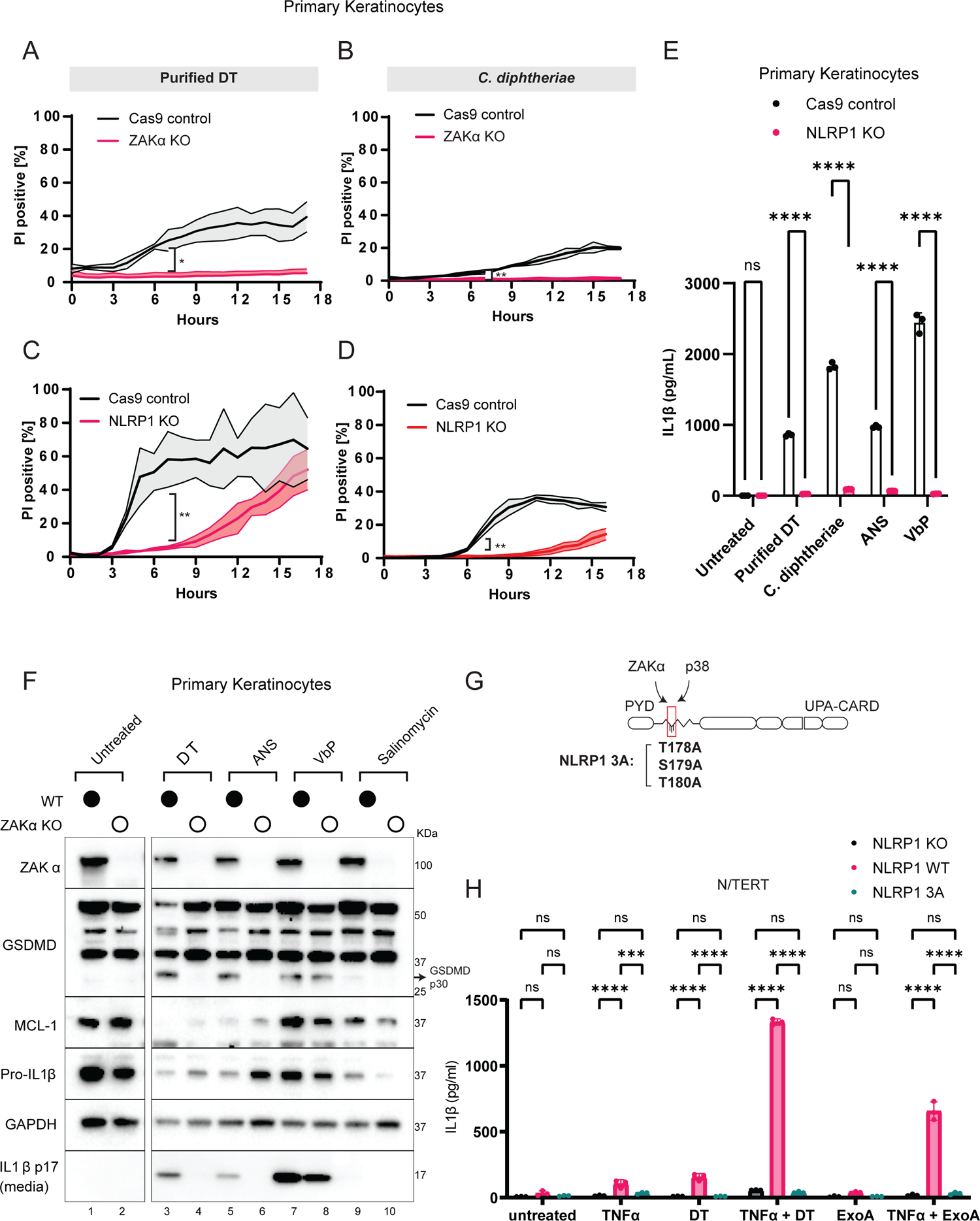
DT and *C. diphtheriae* causes pyroptosis via ZAKɑ and p38-driven NLRP1 phosphorylation. (A) Comparison of PI uptake kinetics between control and ZAKɑ KO primary keratinocytes in response to purified DT and (B) *C. diphtheriae* filtrate. (C) Comparison of PI uptake kinetics between control and NLRP1 KO primary keratinocytes in response to purified DT and (D) *C. diphtheriae* filtrate. Error bars represent three biological replicates, with each drug treatment considered as one replicate. Significance values were calculated from Student’s t test at the 7 hour time point. (E) IL-1β ELISA from primary keratinocyte media 24 hours after the indicated treatment. Significance values were calculated from one way ANOVA with multiple group comparisons. (F) Immunoblot of ZAKɑ, GSDMD, MCL-1 and IL-1β in WT and ZAKɑ KO keratinocyte lysates or media after the indicated treatment. Salinomycin (10 µM) does not activate the NLRP1 inflammasome and was used as a negative control. (G)The location of the critical ZAKɑ and p38 phosphorylation sites on NLRP1 and the construction of the NLRP1 3A mutant. (H) IL-1β ELISA from cell culture media of NLRP1 KO N/TERT cells, or NLRP1 KO N/TERT cells expressing wild-type NLRP1 or NLRP1 3A. Cell culture media were harvested 24 hours after the indicated treatment. Significance values were calculated from one way ANOVA with multiple group comparisons.

### Identification of core RSR signature transcripts in multiple cell types

To gain a more global view on the effects of DT, we performed bulk RNAseq on primary keratinocytes treated with DT for 4 hours in the presence or absence of compound 6p, a more specific and potent ZAKα inhibitor (Yang et al., 2020) (Fig. 4A, Fig. S7A). By analyzing ZAKα-dependent DT-induced transcripts, this experiment serves two purposes: first, to validate the function DT as an RSR inducer using an orthogonal assay, and second, to identify the transcripts controlled by ZAKα-driven RSR while distinguishing them from those mediated by general cytotoxicity. Consistent with biochemical findings shown above, Gene Ontology (GO) analysis of ZAKα-dependent DT-induced transcripts revealed an enrichment for MAPK and IL-18 signaling pathway components (Fig. 4B and 4C, Fig. S7B). The induction of IL-18 response was likely due to paracrine IL-18 signaling originating from a small percentage of cells that had undergone pyroptosis at the time of harvest (Fig. 3A). Interestingly, GO analysis also uncovered a significant overlap between DT and photodynamic therapy (Fig. 4C), a treatment modality for skin hyperplasia that relies on laser irradiation induced cytotoxicity and is mechanistically similar to UV-induced cell death (Choi et al., 2015). As UVB irradiation is known to trigger ZAKα-driven RSR, this finding suggests that different ribotoxic stress agents might induce a common transcriptional program. We thus hypothesized that there exists a common set of RSR signature transcripts that can be induced by all ribotoxic stress agents, regardless of cell type. To investigate this further, we carried out an analogous RNAseq experiment using Cas9 control and ZAKα KO N/TERT cells irradiated with UVB (Fig. S7C-D). Despite the pleiotropic nature of UVB and DT, a significant number of transcripts were induced by both(Fig. S8A-B). These transcripts are enriched for pro-inflammatory cytokines and chemokines (Fig 4D, S8C), stress responsive transcription factors (TFs) (Fig. S8D-E) and ‘early response genes’. Remarkably, many of these shared transcripts are also induced by *L. pneumophila* infection in non-NLRP1 expressing HEK293 cells or murine macrophages (Fontana et al., 2011; Subramanian et al., 2022). Notably, one *L. pneumophila* secreted factor, sidI was recently shown to activate ZAKα-dependent RSR in HEK293 cells (Subramanian et al., 2022). Thus, these data support the notion that ZAKα-driven RSR signaling causes conserved transcriptional changes in diverse cell types, including those which do not express a functional NLRP1 inflammasome. By intersecting our data with those from Subramanian et al (Subramanian et al., 2022), we obtained a more refined cell type-agnostic transcriptional signature that is induced by 5 distinct RSR agents: DT, UVB, ANS, DDB and *L. pneumophila* (Fig. 4E). A stress-responsive TF, ATF3, is induced by all and was shown to mediate cell survival in *L. pneumophila* infection in a sidI-dependent manner. We thus examined whether ATF3 is upstream of the NLRP1 inflammasome or represents a distinct arm of RSR in primary human keratinocytes (Fig. 4E). We first confirmed that ATF3 can also be strongly induced by anisomycin (ANS) in N/TERT cells and primary keratinocytes before the onset of pyroptosis (Fig. S9A and S9C); however, CRISPR/Cas9-mediated deletion of ATF3 did not alter either the kinetics of DT- or ANS-induced pyroptosis as measured by PI uptake, or the level of IL-1β secretion (Fig. 4F, S9B, S9D). These data support a refined and more granular model of RSR: ATF3 and NLRP1 represent two distinct arms of RSR downstream of ZAKα and SAPKs. While ATF3-mediated survival/stress signaling is functional in most cell types, the NLRP1 inflammasome specifically operates in skin keratinocytes and other primary epithelial cells (Fig. 4G and companion manuscript).

**Figure 4.**
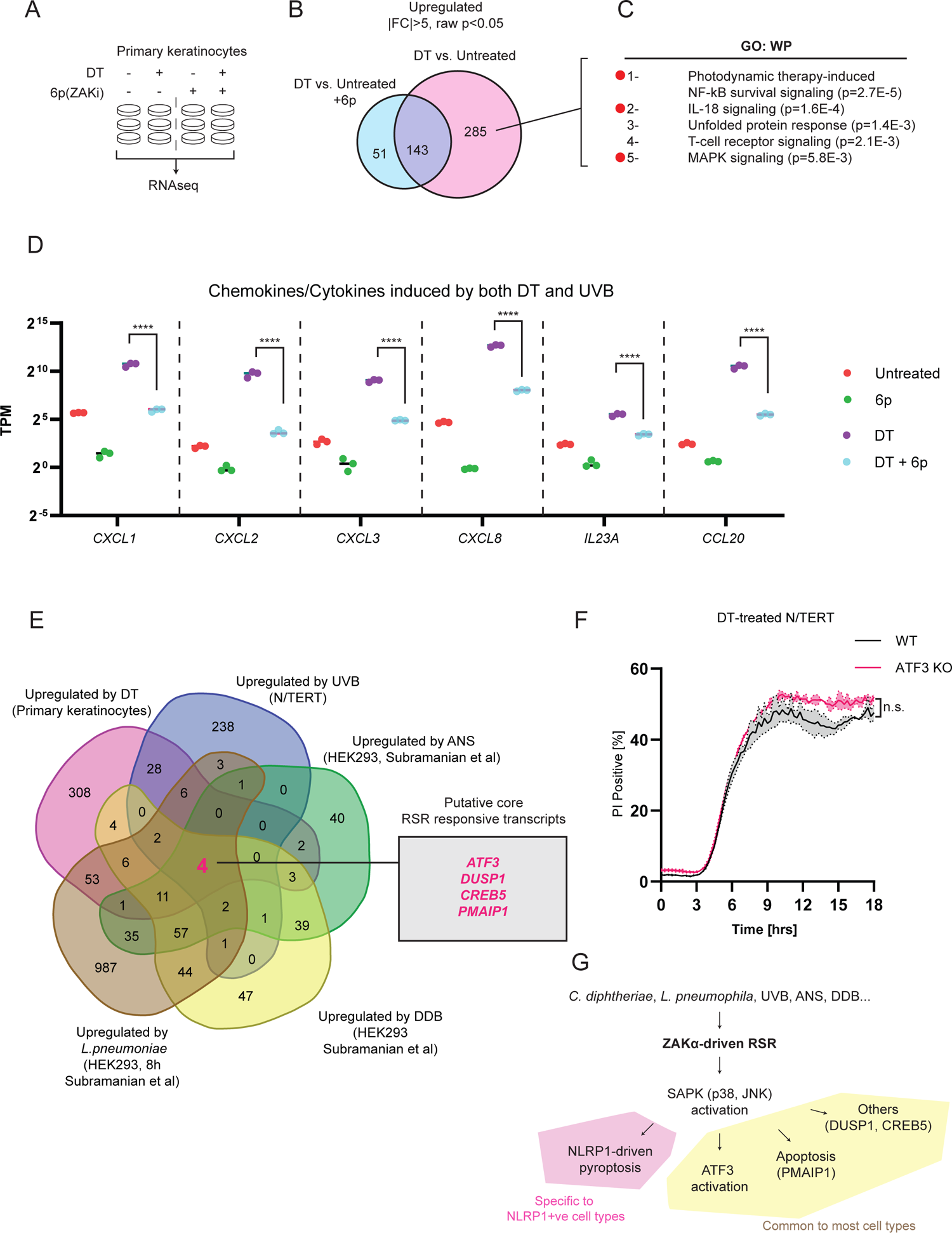
Identification of transcripts that are induced by multiple RSR agents across multiple cell types. (A) Design of RNAseq experiment. (B) Venn diagram showing the identification of 285 ZAKɑ-dependent, DT-induced transcripts in primary keratinocytes. (C) GO analysis of ZAKɑ-dependent DT-induced transcripts. (D) TPM values of select chemokines and cytokines that are upregulated by both DT and UVB in primary keratinocytes described in (C). Significance values were calculated from Student’s t test. (E) Venn diagram demonstrating the overlap of transcriptional changes elicited by the distinct RSR agents. The source cell types are indicated in brackets. FC>2 used a cutoff for datasets published in Subramanian et al. (F) Kinetics of PI uptake of primed wild-type and ATF3 KO N/TERT cells treated with DT. Significance values were calculated from Student’s t test at the 7 hour time point. (G)Proposed model of RSR and its sub-branches.

### ZAKα and p38 inhibitors rescue epithelial integrity in a 3D skin model of cutaneous diphtheria

Next we employed human 3D organotypic skin as an approximate model for cutaneous diphtheria (Fig. 5A). This system offers advantages over murine models, since 1) NLRP1 inflammasome components are not expressed in murine skin (Sand et al., 2018) and muNLRP1 lacks the linker region that harbors the ZAKα and p38 phosphorylation sites (Robinson et al., 2022; Jenster et al., 2023). Furthermore, murine cells in general are resistant to DT entry due to the sequence divergence of the DT receptor, pro-HBEGF (Ivanova et al., 2005). Using H&E staining, we established that recombinant DT causes significant damage to the stratified epidermis in 3D organotypic skin cultures, as evidenced by prominent dyskeratotic and vacuolated keratinocytes with condensed nuclei in basal and spinous layers (Fig. 5B, S10A, yellow arrows). This was accompanied by the weakening of the epidermal-dermal junction and significant detachment of the epidermis from the dermis (Fig. 5B, S10A). These features were also observed in 3D cultures treated with VbP, suggesting that they are likely a consequence of keratinocyte-intrinsic pyroptosis (Fig. 5B). Next we measured the levels of 65 inflammatory cytokines and chemokines in DT- and VbP-induced 3D organotypic skin using Luminex. Remarkably, DT and VbP led to the secretion of a nearly identical set of pro-inflammatory cytokines/chemokines, with IL-18 and IL-1β amongst the most induced Thus, even though VbP and DT have unrelated, non-inflammasome-dependent activities, their effects on 3D skin cultures are dominated by the aftermath of NLRP1 inflammasome-driven pyroptosis. We further hypothesized that pharmacologic inhibition of ZAKα-driven NLRP1 activation could prevent DT-induced epidermal disruption. Indeed, the p38 kinase inhibitor Neflamapimod reduced the extent of cell death and the epidermal detachment in DT-treated 3D skin cultures (Fig. S10A); however, the rescue was partial, in agreement with its effects in 2D keratinocyte culture and the reported role of p38 in NLRP1 activation. Nonetheless, Neflamapimod proved effective at inhibiting DT-induced pro-inflammatory cytokines, including IL-18 and IL-1β (Fig. S10B-D). These results confirm that the keratinocyte-intrinsic activation of the NLRP1 inflammasome, downstream of p38, is largely responsible for DT triggered epidermal disruption and inflammation in reconstructed 3D skin cultures. It is likely other arms of the RSR pathway, such as JNK activation, also contribute to the damage to the epidermis caused by DT.

**Figure 5.**
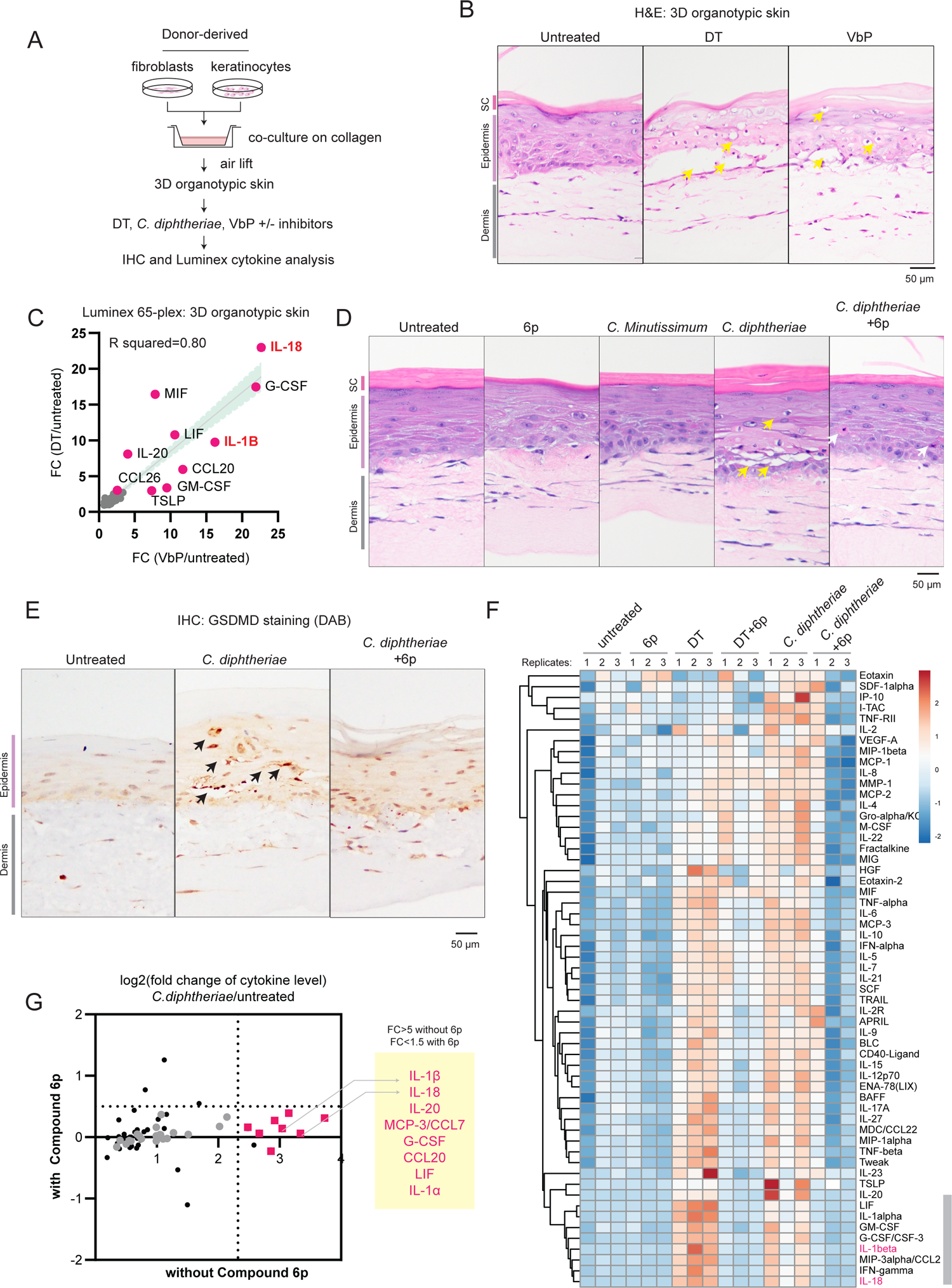
p38 and ZAKα inhibition rescues epidermal integrity by limiting pyroptosis in a model of cutaneous diphtheria. (A) Outline of the 3D human skin model of cutaneous diphtheria. (B) H&E staining demonstrating the histological changes caused by DT and VbP. Yellow arrows indicate dyskeratotic keratinocytes with vacuolated cytoplasm and condensed nuclei. (C) Correlation between the fold change (FC) of 65 cytokines/chemokines after DT or VbP treatment. (D) H&E staining of 3D skin treated with the indicated bacteria filtrate or compound. Yellow arrows indicate dyskeratotic keratinocytes with vacuolated cytoplasm and condensed nuclei. Gray arrows mark putative apoptotic cells with eosin rich cytosol and condensed nuclei. (E) GSDMD p30 staining of 3D skin treated with *C. diphtheriae* filtrate with and without compound 6p. Black arrows indicate membranous staining around foci of epithelial barrier damage. (F) Heatmap demonstrating the relative levels of 65 cytokines/chemokines in triplicate 3D skin cultures treated with the DT, 6p, *C. diphtheriae* filtrates and the indicated combinations. IL-1β and IL-18 are highlighted in red. Row clustering was performed using the Manhattan method. Gray bar indicates the cluster of cytokines that are the most significantly induced by DT and *C. diphtheriae* in a 6p-dependent manner. (G)Log fold change of chemokine/cytokine levels induced by *C. diphtheriae* in 3D skin without 6p (x-axis) and with 6p (y-axis). Gray= chemokine/cytokines that are significantly induced by *C. diphtheriae* without 6p (student’s T test, unpaired). Pink= chemokine/cytokines that are induced by *C. diphtheriae* only in a 6p-dependent manner. Criteria for fold change are indicated.

Finally, we investigated the efficacy of ZAKα inhibition, which blocks the entire RSR pathway, in preventing *C. diphtheria*-induced damage to the 3D skin. Similar to purified DT, filtrate of iron-restricted *C. diphtheria* caused extensive structural damage to the 3D skin, resulting in frequent dyskeratotic keratinocytes and epidermal detachment (Fig. 5D). Furthermore, *C. diphtheria* caused focal loss of adherens junctions and desmosomes in the spinous layer and disorganization of hemidesmosomes in the basal layer, demonstrating that epidermal barrier had been severely compromised (Fig. S11A). This is likely a direct consequence of keratinocyte pyroptosis, as shown by the appearance of membranous staining of cleaved GSDMD p30 in both the basal and spinous layers (Fig. 5E, black arrows, Fig. S11B). These effects were not observed for the related commensal *Corynebacterium* species, *C. Minutissimum* (Fig. S11B). Co-treatment with compound 6p significantly rescued epidermal junctional integrity in *C. diphtheriae*-treated 3D skin as measured by H&E staining (Fig. 5D) and abrogated the GSDMD p30 membranous staining (Fig. 5E, S11A). The cytokines and chemokines induced by *C. diphtheriae* filtrate were similar to those induced by recombinant DT (Fig. 5F). IL-18 and IL-1β were again among the most induced secreted factors by *C. diphtheriae*, further supporting the notion that the NLRP1 inflammasome is the major contributor to the inflammatory aftermath of *C. diphtheriae* infection. Co-incubation with compound 6p not only reduced the levels of IL-18 and IL-1β to near baseline levels (Fig. 5G, S11C-D), but also inhibited other cytokines including IL-20, CCL7, CCL20, IL-1α, G-CSF and LIF (Fig. 5G), which are likely downstream of paracrine IL-18 and IL-1β signaling (Fig. 5C).

Taken together, these results demonstrate that ZAKα inhibition is effective at limiting keratinocyte pyroptosis and inflammatory epidermal damage in a model of *C. diphtheria* skin infection. However, 6p is unable to rescue all aspects of DT-associated toxicity, likely because ZAKα and the NLRP1 inflammasome act downstream of the inhibition of protein synthesis. This might explain why certain DT-induced inflammatory cytokines escape inhibition by Neflamapimod and 6p (Fig. 5F, Fig. S10C), as well as the appearance of eosin-rich, but non-vacuolated keratinocytes in 3D skin treated with *C. diphtheria* and 6p (Fig. 5D, right panel, gray arrows). We postulate that these cells represent keratinocytes undergoing non-pyroptotic cell death, likely as a result of prolonged protein synthesis block.

## DISCUSSION

In summary, we report that multiple bacterial exotoxins that target ribosome elongation factors, EEF1 and EEF2 activate ZAKα-dependent RSR and the NLRP1 inflammasome in primary human epithelial cells. Using cutaneous diphtheria as a model, we demonstrate that one of these toxins, DT, disrupts epidermal barrier integrity via keratinocyte-intrinsic ribotoxic stress and pyroptosis. Pharmacologic inhibition of RSR prevents DT-driven pyroptosis and as a result alleviates *C. diphtheria*-induced skin damage and inflammation. These findings demonstrate that the human NLRP1 inflammasome not only responds to viral pathogens, but also conserved bacterial virulence factors.

Our results have additional implications-although DT is frequently cited as an apoptosis inducing agent, here we show that its predominant effect on human epithelial cells is pyroptosis. As antibody-DT conjugates are currently used to treat certain cancer types (Frankel et al., 2002), the contribution of RSR and the NLRP1 inflammasome in their therapeutic effect warrants further investigation. In addition, given that ZAKα-driven RSR is conserved in rodents (Snieckute et al., 2022; Silva et al., 2022), the immunological consequence of DTA as an *in vivo* cell ablation tool should also be more carefully examined.

In addition, by combining our results with those recently published by (Subramanian et al., 2022), we identified a core set of transcripts that are induced by 5 distinct RSR-inducing agents in multiple cell types. As recent work has implicated RSR in multiple disease states (Vind et al., 2020a), we envision that this ‘transcriptional signature’, upon further refinement, should serve as a useful resource to diagnose if RSR signaling is involved in a particular pathogenesis process, similar to the widely used ‘interferon signature’ (Rice et al., 2017). Our results also imply that ZAKα-dependent RSR has a broader role in innate immunity. We propose that RSR, which encompasses NLRP1-driven pyroptosis in human epithelial cells, represents an example of ‘effector triggered response’ (ETR) (Stuart et al., 2013; Fontana et al., 2011), where an sensor protein (ZAKα, with NLRP1) becomes activated in response to perturbations of an essential cellular process (ribosome elongation) caused by a pathogen virulence factor (DT, exoA and sidI). While RSR- and inflammasome-dependent cytokines could promote bacterial clearance in early stages of diphtheria infection, e.g. by recruiting neutrophils to the infected sites, unchecked NLRP1-driven pyroptosis is most likely a maladaptive and host-detrimental response as the infection progresses. We hypothesize that the loss of epithelial barrier integrity caused by pyroptosis directly promotes the dissemination of DT to internal organs such as the heart at later stages of infection - a frequent fatal complication of diphtheria (Murphy; Sharma et al., 2019; Holmes, 2000). As shown in a companion manuscript, the ZAKα-NLRP1-driven pyroptosis is also a host-detrimental process in *P. aeruginosa* airway infection. In an alternative context, ZAKα-NLRP1-driven ETR might constitute a host-protective response for intracellular pathogens, such as Legionnaires’ disease. This hypothesis awaits additional characterization. It would also be interesting to test if pyroptotic epithelial cells can directly facilitate bacterial growth in their native niche, e.g. by providing additional nutrients or via other means. Notwithstanding, the data shown here should inform future studies that seek to further characterize the arms race between RSR, inflammasome and the microbial ribotoxins, which are abundant in nature (Dmitriev et al., 2020). Finally, in proof-of-concept experiments, we showed that inhibition of ZAKα and p38 signaling effectively blocks DT-induced skin damage in 3D cultures. Although these compounds are unlikely to supplant current treatment and prevention options for diphtheria, ZAKα or NLRP1 inhibitors could find utility for other bacterial infections, such as exoA+ve *P. aeruginosa*, for which effective vaccines and antibiotics are lacking.

## MATERIALS AND METHODS

### Bacterial strains and culture conditions

Two non-toxigenic commensal *Corynebacterium* species, *Corynebacterium striatum* (ATCC 6940) and *Corynebacterium minutissimum* (ATCC 23347) and one toxigenic strain, *Corynebacterium diphtheriae* (ATCC 13812) were cultured in Brain Heart Infusion (BHI) (Sigma-Aldrich, 53286) broth at 37°C overnight in a shaking incubator at 230 rpm. Homemade 611 PGT media with the recipe described in Table S2 was used for quantification of bacterial growth.

### Bacterial growth kinetic and correlation of DT induction

To track bacterial growth under iron starving conditions, and correlate to the amount of DT, C.*Striatum* and C.*Diphtheriae* were incubated at 37°C in a shaking incubator in 611 PGT media containing 0.5mM 2,2’-bipyridyl (Sigma-Aldrich, D2163050). Growth was closely monitored by measuring OD_600_ every hour for 4 hours: the culture was serially diluted 10-fold by sequential transfer of 20 µl into 180 µl of phosphate-buffered saline(PBS) in triplicate wells by a factor of 10^-7^ every hour, for 4 hours. To enumerate bacterial colony forming units (CFU), 5 µl of each dilution was spotted in triplicates on pre-warmed blood agar (Thermofisher Scientific, PP1325P90) and incubated overnight at 37°C. Subsequently, colonies were counted and bacterial concentrations in the original sample estimated. 200 µl of bacteria culture were also harvested every hour for 4 hours, whereby it is centrifuged and supernatant kept to perform immunoblotting in order to correlate the levels of DT induction. 5 ng of purified DT was used as a loading control. The levels of band intensity were used to compare and estimate the amount of DT produced in a given volume of bacteria culture.

### DT induction and harvesting of bacterial sterile filtrates

*C. Diphtheriae* strain was incubated at 37°C in a shaking incubator in BHI broth (Merck, 53286) and was grown to an OD_600_ of 0.4 to 0.6. 2,2’-bipyridyl was added to a final concentration of 0.5mM and incubated for 4 hours to induce diphtheria toxin production under iron starving conditions. The bacteria were then removed by centrifugation at 3000 *xg* for 10 minutes and the supernatant containing DT was harvested and filtered through a 0.2 µm syringe filter. The presence of DT in sterile filtrates is validated through immunoblotting assays with 5 ng of purified DT used as loading control to estimate DT concentration. For cellular and 3D skin culture experiments, unless otherwise indicated, the amount of DT in bacterial filtrate was normalized to be the same as concomitant DT treatment (150 ng/mL).

### Cell culture and chemicals

293Ts (ATCC #CRL-3216) were cultured according to manufacturer’s protocols. Immortalized human keratinocytes (N/TERT herein) were provided by H. Rheinwald (MTA) and cultured in Keratinocyte Serum Free Media (Gibco, 17005042) supplemented with final concentration of 25 mg/l Bovine Pituitary Extract (Gibco, 13028-014), 294.4 ng/l human recombinant EGF (Gibco, 10450-013) and 300 µM of CaCl_2_ (Kanto Chemicals, 07058-00). Primary human keratinocytes were derived from the skin of healthy donors and obtained with informed consent from the Asian Skin Biobank (ASB) (https://www.a-star.edu.sg/sris/technology-platforms/asian-skin-biobank). All cell lines underwent routine mycoplasma testing with MycoGuard (Genecopoeia #LT07-118).

The following drugs and chemicals were used as part of this study: talabostat (VbP, MCE, HY-13233), anisomycin (ANS, MCE, HY18982), neflamapimod (MCE, HY-10328), M443 (MCE, HY-112274), harringtonine (HTN, MCE, HYN0862), puromycin (Sigma, P9620), emricasan (MCE, HY-10396), aplidine (MCE, HY-16050), vemurafenib (MCE, HY-12057), salinomycin (MCE, HY-15597). Compound 6p is a kind gift from X. Lu (Jinan University, China). The following recombinant protein were used in this study: TNFα (R&D Systems, 210-TA), diphtheria toxin (DT, Sigma, D0564), Exotoxin A (ExoA, Sigma, P0184), Streptolysin O (SLO, Sigma, SAE0086). Recombinant LFn-DT and PA were a gift from F. Schmidt (University of Bonn, Germany).

### Organotypic 3D skin culture

Organotypic cultures were generated by adapting a previously described protocol (Arnette et al., 2016). Briefly, 2 ml of collagen I (4 mg/ml) (Corning, #354249) mixed with 7.5×10^5^ human fibroblasts were allowed to polymerize over 1 ml of acellular collagen I in six-well culture inserts (Falcon, #353102) placed in six-well deep well plates (Falcon, #355467). After 24 hours, 1×10^6^ primary human keratinocytes were seeded into the inserts and kept submerged in a 3:1 DMEM (Hyclone, #SH30243.01) and F12 (Gibco, #31765035) mixture with 10% FBS (Hyclone, #SV30160.03), 100 U/ml of penicillin–streptomycin (Gibco, #15140122), 10 µM Y-27632 (Tocris, #1254), 10 ng/ml of EGF (Sigma-Aldrich, #E9644), 100 pM cholera toxin (Enzo, #BML-G117-001), 0.4 µg/ml of hydrocortisone (Sigma-Aldrich, #H0888), 0.0243 mg/ml adenine (Sigma-Aldrich, #A2786), 5 µg/ml of insulin (Sigma-Aldrich, #I2643), 5 µg/ml of transferrin (Sigma-Aldrich, #T2036), and 2 nM 3,3′,5′-triiodo-L-thyronine (Sigma-Aldrich, #T6397). After another 24 hours, the organotypic cultures were then raised at the air–liquid interface and fed with the submerged media (without Y-27632 and EGF) below the insert to induce epidermal differentiation. The air-lifting medium was replaced every 2 days and treatments began 10-14 days after airlifting. Organotypic cultures were then harvested 24 hours after treatment and formalin fixed for 24 hours. Fixed tissues were then embedded into wax for histological purposes before being cut and stained using a standard H&E protocol, the previously described method for GSDMD staining (Robinson et al., 2020) plectin (BD Transduction Labs, #611348) or plakoglobin (Progen #61005S) using standard DAB staining method.

### Cytokine analysis

Human IL-1β enzyme linked immunosorbent assay (ELISA) kit (BD, #557953) was used according to the manufacturer’s protocol. Culture supernatants of 3D skin were also collected and sent for Luminex analysis using the ProcartaPlex, Human Customized 65-plex Panel (Thermo Fisher Scientific) to measure the following targets: APRIL; BAFF; BLC; CD30; CD40L; ENA-78; Eotaxin; Eotaxin-2; Eotaxin-3; FGF-2; Fractalkine; G-CSF; GM-CSF; Gro α; HGF; IFN-α; IFN-γ; IL-10; IL-12p70; IL-13; IL-15; IL-16; IL-17α; IL-18; IL-1α; IL-1β; IL-2; IL-20; IL-21; IL-22; IL-23; IL-27; IL-2 R; IL-3; IL-31; IL-4; IL-5; IL-6; IL-7; IL-8; IL-9; IP-10; I-TAC; LIF; MCP-1; MCP-2; MCP-3; M-CSF; MDC; MIF; MIG; MIP1α; MIP-1β; MIP-3 A; MMP-1; NGF β; SCF; SDF-1 α; TNF-β; TNF-α; TNF-R2; TRAIL; TSLP; TWEAK; VEGF-α.

### RNAseq sample preparation and analysis

Primary human keratinocytes or N/TERT cells were grown to 80% confluence in a 6 well plate before performing treatments. Cells were treated with stated treatments and harvested after 5 hours. Total RNA was isolated from each treatment using RNAeasy mini kit (Qiagen, cat. # 74004). The quantity and quality of each RNA sample was then assayed using the NanoDrop (Thermo Scientific) but also by running the RNA on an agarose gel to check for RNA degradation. RNA sequencing was performed at Macrogen Asia using the Novaseq 6000 platform. Library construction and sequencing followed the standard sequencing protocols and were performed by Macrogen Asia. Preprocessing and analysis was also performed by Macrogen Asia, in brief reads were mapped to reference genome (*homo sapiens*, GRCh38) with HISAT2 and transcript assembled by StringTie with aligned reads. Expression profiles were represented as read count and normalization values which were calculated as TPM (Transcripts Per Kilobase Million) for each sample, based on transcript length and depth of coverage. Using pairwise comparisons, differentially expressed transcripts were calculated using an R package (DEseq2). The DEseq2 analysis was performed on read counts of expressed genes and for each gene the *P-*value and fold change was calculated per comparison pair.

### Plasmids and preparation of lentiviral stocks

293T-ASC-GFP-NLRP1, N/TERT NLRP1 KO + NLRP1^DR^-GFP, N/TERT ZAKα KO, N/TERT p38 dKO cells were previously described (Robinson et al., 2022). All expression plasmids for transient expression were cloned into the pCS2+ vector backbone and cloned using InFusion HD (Clontech). Constitutive lentiviral expression was performed using pCDH vector constructs (System Biosciences) and packaged using third-generation packaging plasmids. SidI plasmids for expression in 293Ts were gifts from S. Mukherjee (University of California San Francisco, USA).

### CRISPR-Cas9 knockout

NLRP1 KO, MAP3K20 (ZAKα) KO, and MAPK14/MAPK11 (p38α/β) dKO N/TERT keratinocytes were made and described in detail previously (Robinson et al., 2022). MAP3K20 (ZAKα) KO human primary keratinocytes were generated using lentiviral Cas9 and guide RNA plasmid (LentiCRISPR-V2, Addgene plasmid #52961) using the following guides: sg1 (TGTATGGTTATGGAACCGAG), sg4 (TGCATGGACGGAAGACGATG). NLRP1 KO human primary keratinocytes were generated using lentiviral transduction with the following guide: sg1 (GATAGCCCGAGTGCATCGG). ATF3 KO N/TERT and human primary keratinocytes were generated using RNP nucleofection of both guides: CTTTTGTGATGGACACCCCG, TAACCTGACGCCCTTTGTCA. DPH1 KO N/TERTs were generated using RNP nucleofection of both guides: GATGGGTGACGTGACCTACG, GCTGACTTCTTGGTGCACTA. Knockout efficiency was tested by immunoblot. Alternatively, Sanger sequencing of genomic DNA and overall editing efficiency were determined using the Synthego ICE tool (Synthego Performance Analysis, ICE Analysis. 2019. v2.0. Synthego, https://ice.synthego.com/<u>#/</u>).

### Harringtonine ribosome runoff assay

Human primary keratinocytes were seeded at a cell density of 80,000 cells/well in a 24 well plate. The next day, cells were pre-treated with 10µM of emricasan (MCE, HY-10396) for 30 minutes followed by treatment with either BHI only, *C. Diphtheriae* or *C. Striatum* for 6.5 hours. Recombinant DT was added to the cells during the final 3 hours. The respective samples were treated with 2µg/ml of harringtonine (MCE, HY-10396) at staggered timepoints of 5 min, 1 min and 0 min (equivalent to mock untreated). Cells were then pulsed with a final concentration of 10 µg/ml of puromycin at the same time in all wells for 10 mins. Following puromycin treatment, the supernatant was discarded and cells were lysed directly in 1x Laemmli buffer. Immunoblotting of samples using an anti-puromycin antibody was done to measure the amount of nascent peptides with puromycin incorporated, which reflects the rate of elongation after DT treatment.

### Propidium iodide inclusion assay

N/TERT cells or human primary keratinocytes of various genotypes were seeded at a cell density of 10,000 cells/well in a black 96-well plate (PerkinElmer, CellCarrier-96 Ultra, #6055300) or a cell density of 80,000 cells/well in black 24-well plates (Cellvis, P24-1.5P). The next day, cells were treated with chemicals and stained with 0.5 µg/ml of propidium iodide (PI, Abcam #ab14083) before observing the cells on a high content screening microscope (Perkin Elmer Operetta CLS imaging system, NTU Optical Bio-Imaging Centre in Nanyang Technological University, Singapore) over 18 hours, capturing brightfield and fluorescent images every 15 mins. For five fields of view per well in 96-well format with three wells per treatment, or 14 fields of view per well in 24-well format, the ratio of PI positive cells over normal cells was calculated. The number of live cells per field was counted using digital phase contrast images, which can identify cell borders, whereas the number of PI stained regions identified through the PI channel (536 nm/617 nm) was counted as PI + cells.

### ASC-GFP speck quantification

Images of ASC-GFP specks were acquired in 20 random fields in 4′,6-diaminidino-2-phenylindole (DAPI, 358 nm/461 nm) and GFP (469 nm/525 nm) channels under 20x magnification using the Operetta CLS imaging microscope. The number of DAPI-stained nuclei and ASC specks in the GFP channel was counted using the Operetta CLS analysis system.

### Immunoblotting

For SDS-PAGE using whole cell lysates, cells were resuspended in tris-buffered saline 1% NP-40 with protease inhibitors (Thermo Scientific, #78430). Protein concentration was determined using the Bradford assay (Thermo Scientific, #23200) and 20 µg of protein loaded. To visualize cleaved GSDMD, cell debris floaters were separated after collection of media supernatants post treatment and lysed directly with 1x Laemmli Buffer (30ul of 1xLB for 200,000 cells). Attached cells on the wells were lysed directly with 1x Laemmli Buffer (100ul of 1xLB for 200,000 cells), sonicated and boiled. Lysates from the floater fraction and the attached cell fraction were loaded into each gel lane in a 1:1 ratio (10ul floaters, 10ul attached cells’ lysate). For analysis of IL-1β in the media by immunoblotting, samples were concentrated using filtered centrifugation (Merck, Amicon Ultra, #UFC500396). Protein samples were run using immunoblotting, and then visualized using a ChemiDoc Imaging system (Bio-Rad).

PhosTag SDS-PAGE was carried out using homemade 10% SDS-PAGE gel, with addition of Phos-tag Acrylamide (Wako Chemicals, AAL-107) to a final concentration of 30 µM and manganese chloride(II) (Sigma-Aldrich, #63535) to 60 µM. Cells were directly harvested using 1x Laemmli buffer, lysed with an ultrasonicator, and loaded into the Phos-tag gel to run. Once the run was completed, the polyacrylamide gel was washed in transfer buffer with 10 mM EDTA twice, subsequently washed once without EDTA, blotted onto 0.45 µm PVDF membranes (Bio-rad), blocked with 3% milk, and incubated with primary and corresponding secondary antibodies. All primary and secondary antibodies used in this study are mentioned in Table S1 below.

---

**Table S1.**
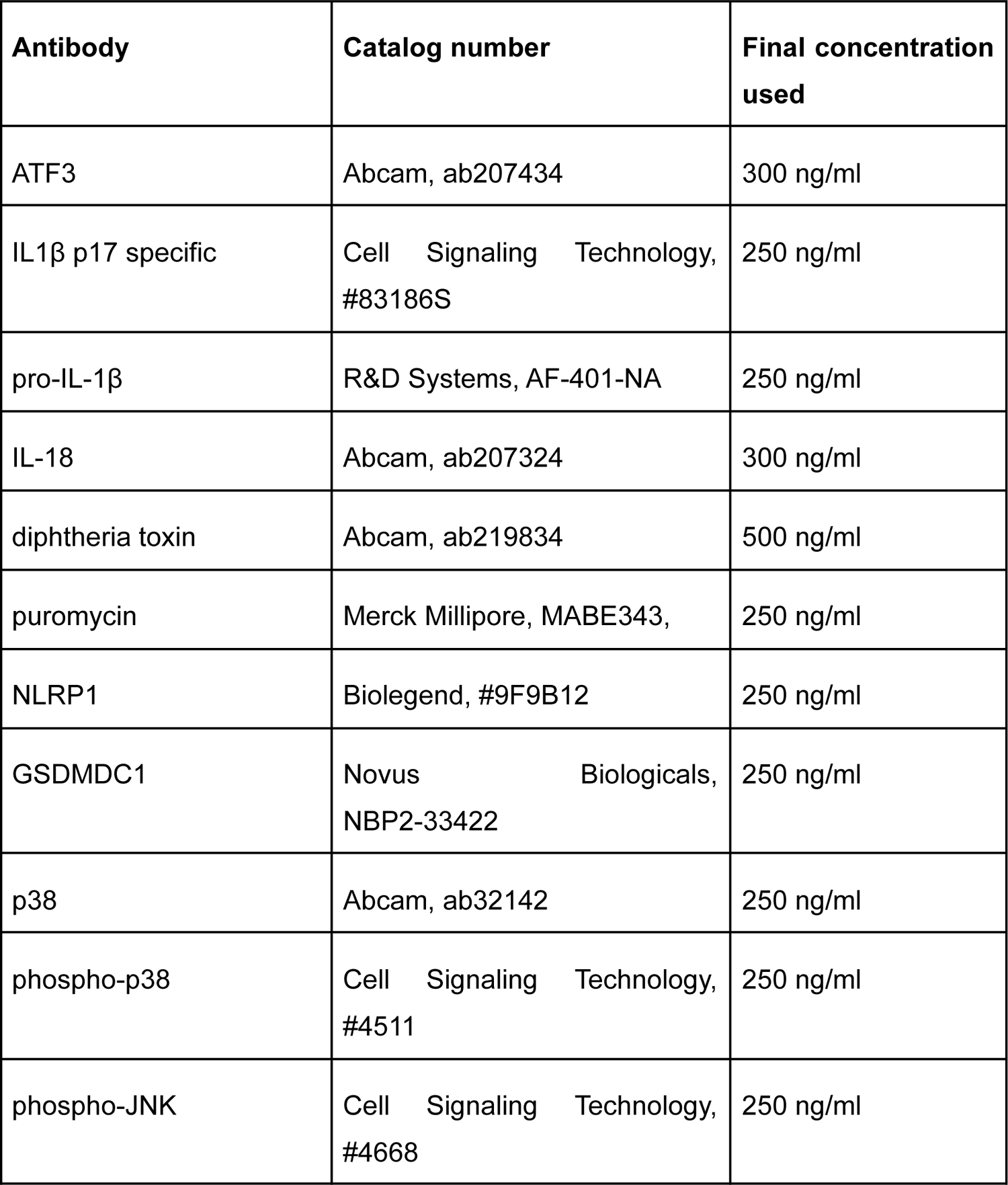
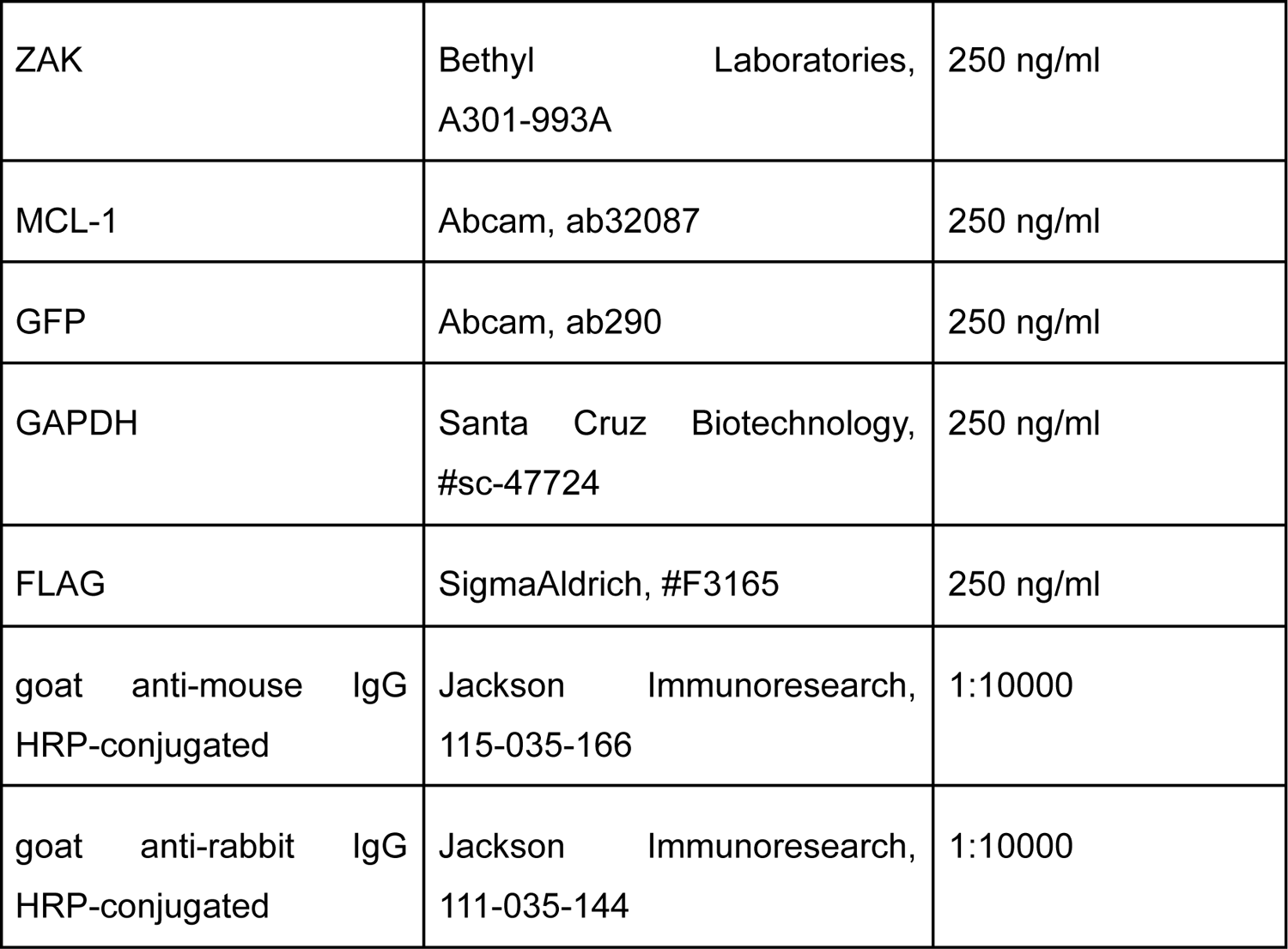
Antibodies used in this study.

| **Antibody** | **Catalog number** | **Final concentration used** |
| --- | --- | --- |
| ATF3 | Abcam, ab207434 | 300 ng/ml |
| IL1β p17 specific | Cell Signaling Technology, #83186S | 250 ng/ml |
| pro-IL-1β | R&D Systems, AF-401-NA | 250 ng/ml |
| IL-18 | Abcam, ab207324 | 300 ng/ml |
| diphtheria toxin | Abcam, ab219834 | 500 ng/ml |
| puromycin | Merck Millipore, MABE343, | 250 ng/ml |
| NLRP1 | Biolegend, #9F9B12 | 250 ng/ml |
| GSDMDC1 | Novus Biologicals, NBP2-33422 | 250 ng/ml |
| p38 | Abcam, ab32142 | 250 ng/ml |
| phospho-p38 | Cell Signaling Technology, #4511 | 250 ng/ml |
| phospho-JNK | Cell Signaling Technology, #4668 | 250 ng/ml |
| ZAK | Bethyl Laboratories,<br>A301-993A | 250 ng/ml |
| MCL-1 | Abcam, ab32087 | 250 ng/ml |
| GFP | Abcam, ab290 | 250 ng/ml |
| GAPDH | Santa Cruz Biotechnology,<br>#sc-47724 | 250 ng/ml |
| FLAG | SigmaAldrich, #F3165 | 250 ng/ml |
| goat anti-mouse IgG<br>HRP-conjugated | Jackson ImmunoResearch,<br>115-035-166 | 1:10000 |
| goat anti-rabbit IgG<br>HRP-conjugated | Jackson ImmunoResearch,<br>111-035-144 | 1:10000 |

**Table S2.**
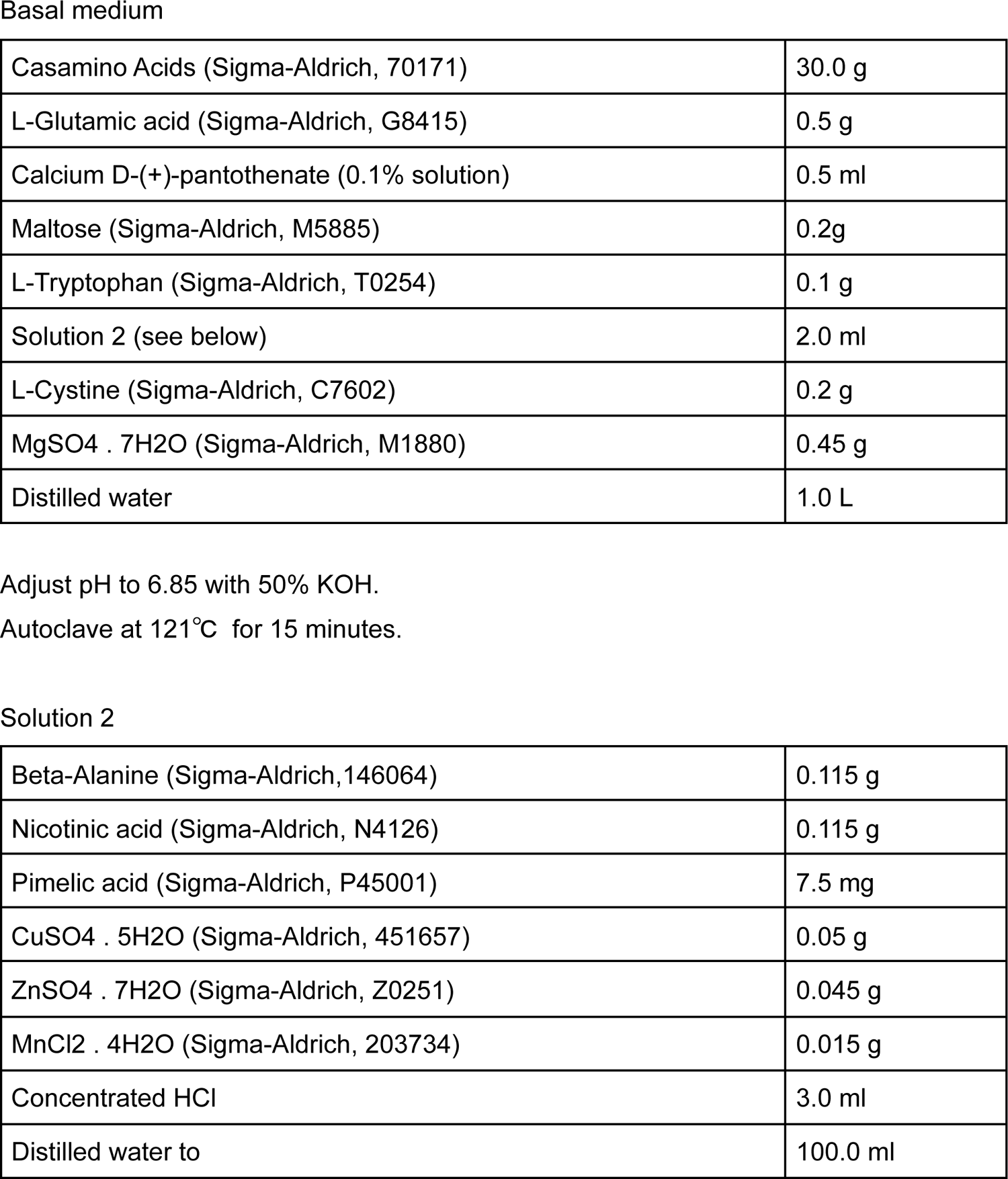
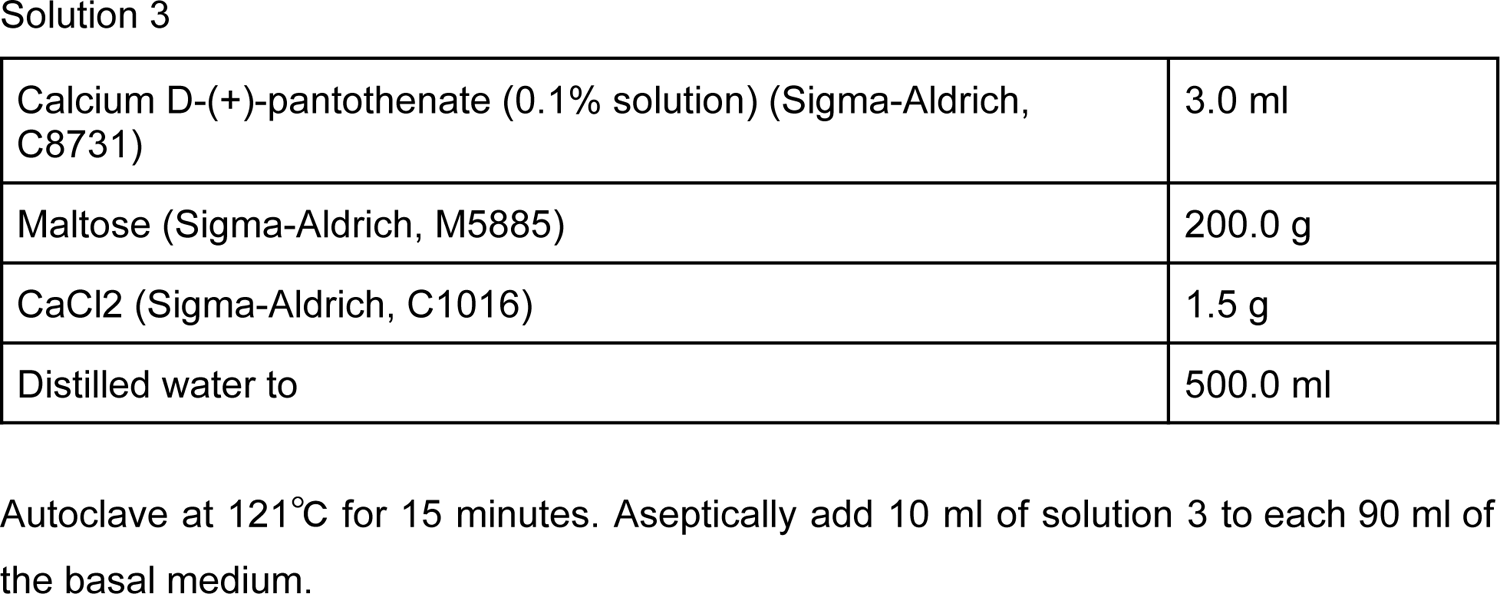
Composition of the 611 PGT Medium (adapted from ATCC)

|  |  |
| --- | --- |
| Casamino Acids (Sigma-Aldrich, 70171) | 30.0 g |
| L-Glutamic acid (Sigma-Aldrich, G8415) | 0.5 g |
| Calcium D-(+)-pantothenate (0.1% solution) | 0.5 ml |
| Maltose (Sigma-Aldrich, M5885) | 0.2g |
| L-Tryptophan (Sigma-Aldrich, T0254) | 0.1 g |
| Solution 2 (see below) | 2.0 ml |
| L-Cystine (Sigma-Aldrich, C7602) | 0.2 g |
| MgSO<sub>4</sub> · 7H<sub>2</sub>O (Sigma-Aldrich, M1880) | 0.45 g |
| Distilled water | 1.0 L |
| Beta-Alanine (Sigma-Aldrich, 146064) | 0.115 g |
| Nicotinic acid (Sigma-Aldrich, N4126) | 0.115 g |
| Pimelic acid (Sigma-Aldrich, P45001) | 7.5 mg |
| CuSO<sub>4</sub> · 5H<sub>2</sub>O (Sigma-Aldrich, 451657) | 0.05 g |
| ZnSO<sub>4</sub> · 7H<sub>2</sub>O (Sigma-Aldrich, Z0251) | 0.045 g |
| MnCl<sub>2</sub> · 4H<sub>2</sub>O (Sigma-Aldrich, 203734) | 0.015 g |
| Concentrated HCl | 3.0 ml |
| Distilled water to | 100.0 ml |
| Solution 3 |  |
| Calcium D-(+)-pantothenate (0.1% solution) (Sigma-Aldrich, C8731) | 3.0 ml |
| Maltose (Sigma-Aldrich, M5885) | 200.0 g |
| CaCl<sub>2</sub> (Sigma-Aldrich, C1016) | 1.5 g |
| Distilled water to | 500.0 ml |

Autoclave at 121°C for 15 minutes. Aseptically add 10 ml of solution 3 to each 90 ml of the basal medium.

## ACKNOWLEDGEMENTS

The authors are grateful for useful discussion and scientific advice from all members of the Zhong lab. We are especially grateful for Dr. Vijaya Chandra Shree Harsha (A*SRL) for his help with *Corynebacterium* culture. F.L.Z. would like to thank Prof. Veit Hornung (Gene Center, Ludwig-Maximilians-University Munich), Prof. Simon Bekker-Jensen (University of Copenhagen), Prof. Lena Ho (Duke-NUS), Prof. Wu Bin (NTU) and Prof. Seth Masters (WEHI, Australia) for scientific advice and guidance. Work from F.L.Z.’s lab is funded by the National Research Foundation Fellowship, Singapore (NRF-NRFF11-2019-0006) and Nanyang Assistant Professorship (NAP).. Work from J.C.’s lab is supported by funding from Agency for Science, Technology and Research (A*STAR) and A*STAR-EDB-NRF IAF-PP grants – H17/01/a0/004 ‘Skin Research Institute of Singapore’ (K.S.R., C.B., J.E.A.C. and F.L.Z.). and H22J1a0040 ‘Asian Skin Microbiome Program 2.0’ (K-C.T. and J.E.A.C.).

## Notes

### Competing Interest Statement

The authors have declared no competing interest.

