## Supplementary FigS1-S11 and legends for "*Corynebacterium diphtheriae* causes keratinocyte-intrinsic ribotoxic stress and NLRP1 inflammasome activation in a model of cutaneous diphtheria"

#### SUPPLEMENTAL FIGURE LEGENDS

##### Figure S1. Additional data demonstrating the DT activates the NLRP1 inflammasome

(A) Immunoblot of inflammasome components NLRP1, GSDMD and IL-1β in N/TERT with and without TNFα priming (25 ng/mL, 8 hours). Streptolysin O (SLO, 1 μg/ml) was included as additional negative control.

(B) IL-1β ELISA of culture media from N/TERT cells treated with the indicated recombinant proteins or compounds. LFn (150 ng/mL): Lethal Factor from *B.anthracis* (aa 34-288). PA (300 ng/mL): Protective Antigen. MxiH: *Shigella flexneri* type 3 secretion needle protein. All cells were primed with TNFα. Significance values were calculated from one way ANOVA with multiple group comparisons.

(C) Kinetics of PI uptake in TNFα-primed N/TERT cells treated with DT or LFn-DTA+PA.

**Figure S2. DDB/Aplidine and sidI activate the inflammasome**

(A) Kinetics of PI uptake in unprimed N/TERT cells treated with DDB/Aplidine. Significance values were calculated from Student's t test at 4 hours. Data represents one of two biological replicates.

(B) IL-1β ELISA from the N/TERT culture media 18 hours after the indicated drug treatment. Media were harvested 18 hours after treatment. Significance values were calculated from one way ANOVA with multiple group comparisons. Data represents one of two biological replicates.

(C) The percentage of cells with ASC-GFP specks among 293T-ASC-GFP-NLRP1 cells transfected with the indicated plasmids. Cells were fixed 24 hours after transfection. ASC-GFP specks were visualized using GFP epifluorescence and normalized to the total number of cells per field of view using DAPI nuclear counterstain.

(D) Immunoblot of overexpressed 3xFLAG-sidI and R453P glycolysase-defective mutant. Note that the level of wild-type sidI is much lower than the R453P mutant due to its strong inhibitory effect on translation, as reported previously (Subramanian et al. 2022).

**Figure S3. The effect of iron depletion on DT**

(A) Anti-DT immunoblot from the indicated bacterial broth. 2,2'-dipyridyl was added 4 hours before harvest. The broth volume and bacterial counts were normalized by optical density measurement (OD600).

(B) Anti-DT immunoblot from the indicated bacterial strains. All bacteria were treated with 2,2'-dipyridyl for 4 hours.

(C) Correlation between OD600 and colony forming units (CFUs) of *C. diphtheriae* culture through 8 hours, in the presence of 2,2'-dipyridyl. The linear fit curve was used to estimate the amount of DT produced per million bacteria.

(D) Immunoblot of inflammasome activator markers GSDMD and IL-1β in primary keratinocytes treated with the indicated bacterial filtrate.

**Figure S4. DPH1 KO eliminates DT-induced pyroptosis.**

(A) ICE analysis of DPH1 KO N/TERT cells.

(B) IL-1β ELISA from TNFα-primed wild-type or DPH1 KO N/TERT culture media. Media were harvested 18 hours after treatment. Significance values were calculated from one way ANOVA with multiple group comparisons, from 3 biological replicates.

**Figure S5. Additional data demonstrating that ZAKα and NLRP1 are genetically required for DT-induced pyroptosis**

(A) Immunoblot of endogenous ZAKα in control and ZAKα KO primary human keratinocytes.

(B) Kinetics of PI uptake for untreated control and ZAKα KO primary keratinocytes.

(C) Kinetics of PI uptake for untreated control and NLRP1 KO primary keratinocytes.

(D) Immunoblot of inflammasome activator markers from the lysates and media of wild-type and NLRP1 KO primary keratinocytes. Cells were harvested 24 hours after the indicated treatment.

(E) Immunoblot of inflammasome activation markers from the lysates and media of wild-type primary keratinocytes in the presence of M443. Cells were harvested 24 hours after the indicated treatment.

**Figure S6. The role of p38 and NLRP1 linker phosphorylation in DT- and exoA-induced pyroptosis**

(A) IL-1β ELISA of control, ZAKα KO and p38 dKO N/TERT cells treated with DT or exoA. TNFα was added 24 hours prior to DT or exoA. Significance values were calculated from one way ANOVA with multiple group comparisons, from 3 biological replicates.

(B) Kinetics of PI uptake of TNFα-primed N/TERT cells treated with DT and the indicated inhibitors (1 μM). Significance values were calculated from Student's t test at the 7 hour time point.

(C) IL-1β ELISA of control, ZAKα KO and p38 dKO N/TERT cells treated with DT or exoA, and the indicated inhibitors. TNFα was added 24 hours prior to DT or exoA. Significance values were calculated from one way ANOVA with multiple group comparisons, from 3 biological replicates.

(D) Kinetics of PI uptake of DT-treated TNFα-primed NLRP1 KO N/TERT cells rescued with wild-type NLRP1 or NLRP1 3A mutant.

(E) Kinetics of PI uptake of exoA-treated TNFα-primed NLRP1 KO N/TERT cells rescued with wild-type NLRP1 or NLRP1 3A mutant. Significance values were calculated from Student's t test at the 7 hour time point.

**Figure S7. Additional RNAseq experiments to identify 'core' RSR-inducible transcripts**

(A) Percentage of IL-1β inhibition by the indicated titrations of ZAKα inhibitors. IL-1β ELISA was performed using TNFα-primed N/TERT media treated with ANS and the indicated inhibitors 24 hours after treatment. The percentage of inhibition was normalized to the treatment condition where no inhibitor was added. Significance values were calculated from Student's t test.

(B) Heatmap and unsupervised clustering of differentially expressed genes (DEGs) induced by DT in the presence or absence of 6p in primary keratinocytes.

(C) Design of RNAseq experiments to identify UVB-regulated genes that are also sensitive to ZAKα.

(D) Heatmap and hierarchical clustering of DEGs induced by UVB in wild-type and ZAKα KO N/TERT cells.

**Figure S8. Further analysis of transcripts that are induced by multiple RSR agents**

(A) Venn diagram showing that 212 transcripts are induced by UVB in a ZAKα dependent manner in N/TERT cells.

(B) Venn diagram showing that 40 transcripts are upregulated by both UVB and DT.

(C) TPM transcript levels of select chemokine and cytokine transcripts that are induced by both DT and UVB in mock and UVB-irradiated N/TERT cells.

(D), (E) TPM transcript levels of transcription factors that are induced by both DT and UVB. Significance values were calculated from Student's t test.

**Figure S9. Further data demonstrating that the NLRP1 inflammasome is not downstream of ATF3**

(A) ATF3 immunoblot in control and ATF3 KO N/TERT cells. Lysates were harvested 3 hours after ANS.

(B) IL-1β ELISA of control and ATF3 KO primed N/TERT cells after the drug/toxin treatments. Significance values were calculated from one way ANOVA from two biological replicates.

(C) ATF3 immunoblot in control and ATF3 KO primary keratinocytes. Lysates were harvested 3 hours after ANS

(D) Immunoblot of GSDMD and IL-1β comparing the inflammasome response of Cas9 control and ATF3 KO keratinocytes.

**Figure S10. Further analysis of the effects of *C. diphtheriae* on 3D human skin**

(A) H&E staining of 3D skin treated with DT and Neflamapimod.

(B) Principal component analysis of the cytokine and chemokines levels (Luminex 65-plex) in 3D skin culture media after the indicated treatments. Numbers indicate biological replicates

(C) Heatmap showing the levels of 65 cytokines/chemokines in triplicates 3D skin samples treated with the VbP, DT, Neflammapimod (0.5 μM) or Neflammapimod+DT. IL-1β and IL-18 are highlighted in red.

(D) Significantly upregulated cytokines/chemokines in DT-treated 3D skin culture relative to DT+Neflamapimod treated 3D skin. Log10(q values) and log2(fold change) were calculated from a multiparametric t-test, with a false discovery rate of 5%.

**Figure S11. The effect of compound 6p on *C. diphtheriae*-induced skin damage in 3D skin cultures.**

(A) Immunohistochemistry (IHC) staining of plakoglobin (desmosome and adherens junction marker) and plectin (hemidesmosome marker) in 3D skin sections treated with the indicated conditions. Black arrows indicate foci of epidermal damage with loss of plakoglobin staining. Brackets indicate areas of epidermal-dermal junction with disorganized plectin staining.

(B) Additional GSDMD IHC staining of 3D skin sections with control treatment conditions.

(C) IL-1β ELISA of 3D skin culture media after the indicated treatments. Significance values were calculated from one way ANOVA from three biological replicates shown in Fig. 5.

(D) Western blot of IL-18 and IL-1β p17 in the cultured media of 3D skin after the indicated treatment.

### Figure S1

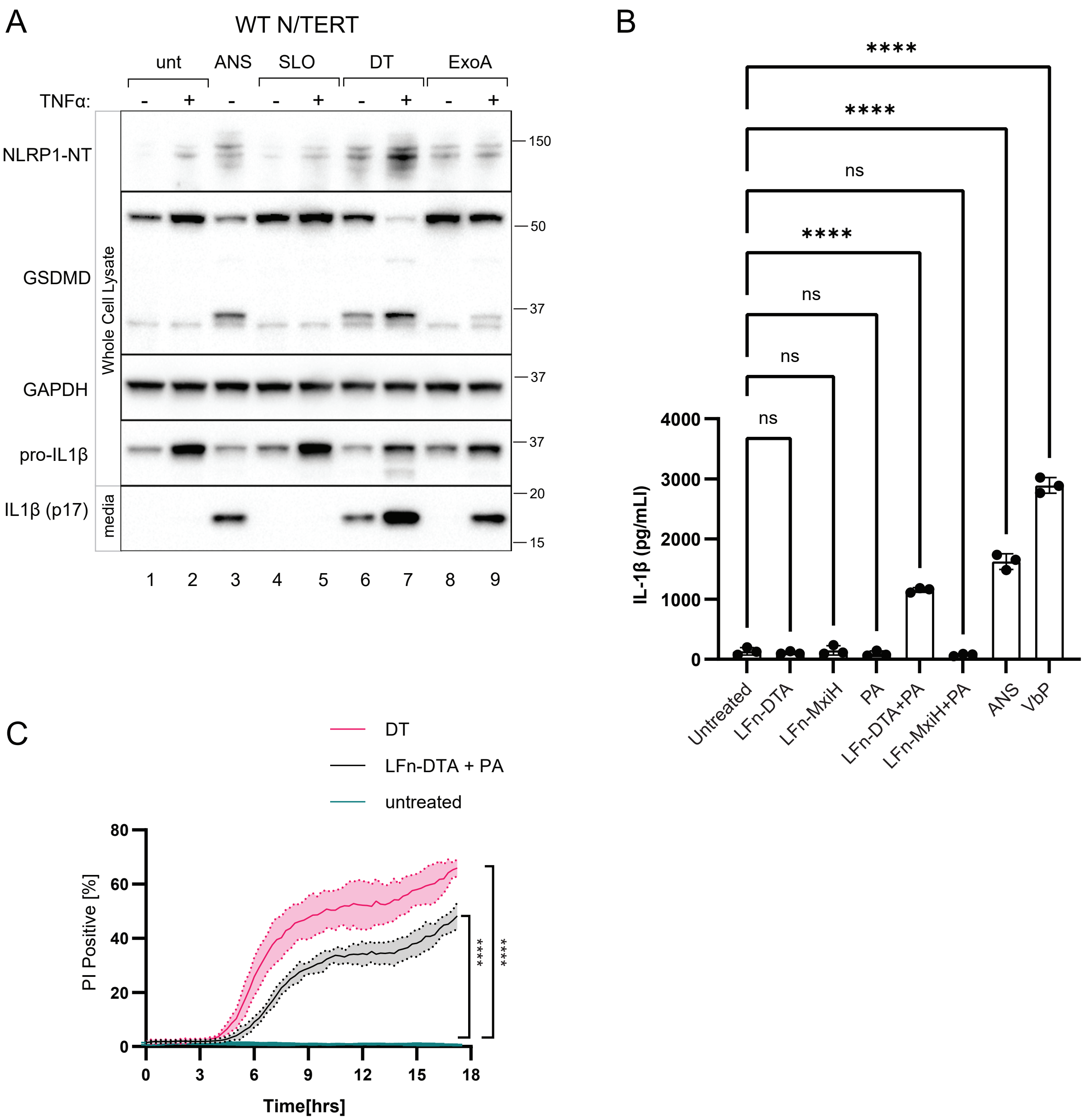
A
WT N/TERT
unt
ANS
SLO
DT
ExoA
TNFα:
NLRP1-NT
GSDMD
GAPDH
pro-IL1β
IL1β (p17)
Whole Cell Lysate
media
150
50
37
37
37
20
15
1 2 3 4 5 6 7 8 9
B
****
****
ns
****
ns
ns
ns
IL-1β (pg/mLl)
4000
3000
2000
1000
0
Untreated
LFn-DTA
LFn-MxiH
PA
LFn-DTA+PA
LFn-MxiH+PA
ANS
VbP
C
DT
LFn-DTA + PA
untreated
PI Positive [%]
80
60
40
20
0
0 3 6 9 12 15 18
Time[hrs]
****
****

### Figure S2

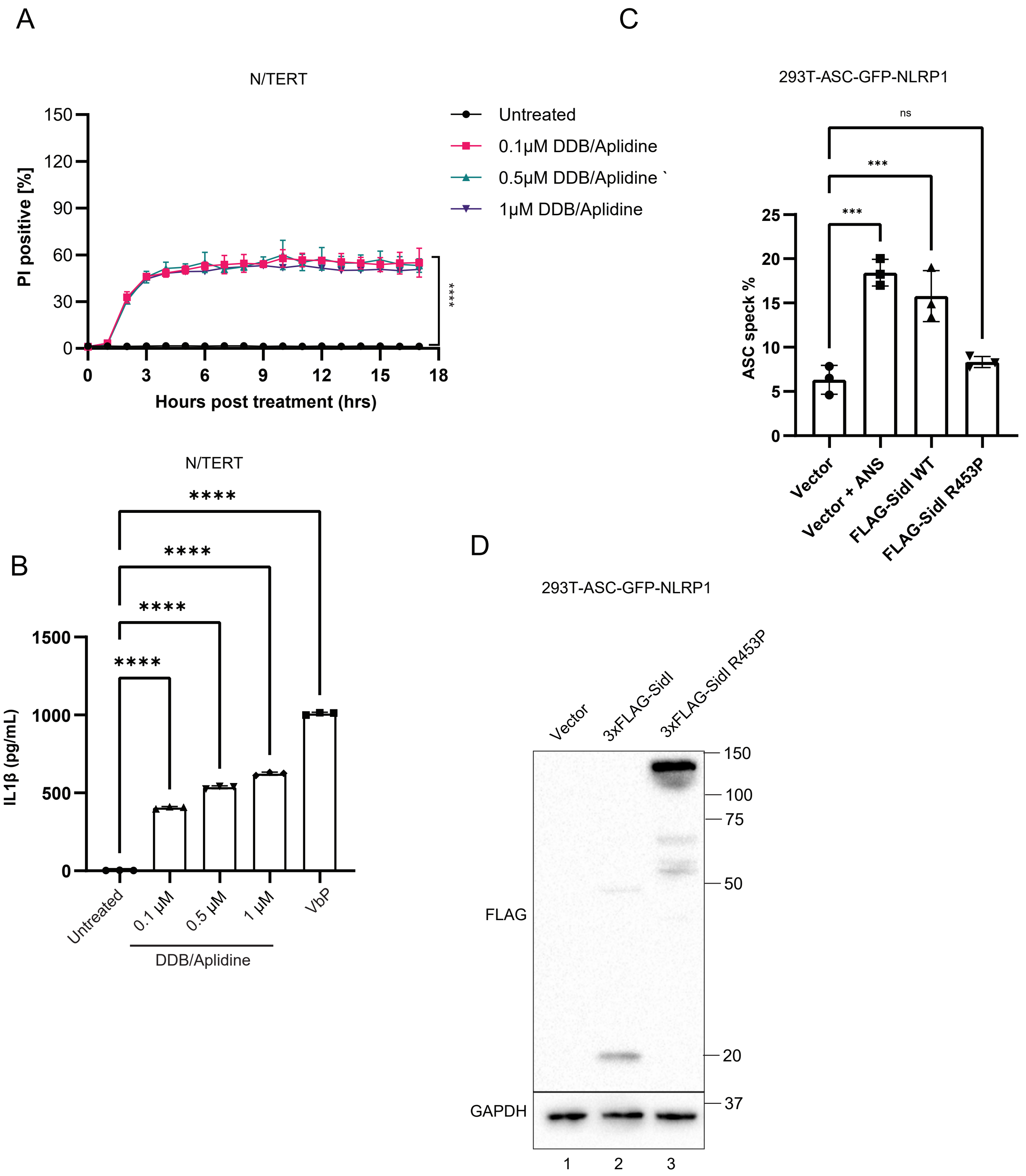
A
N/TERT
PI positive [%]
Hours post treatment (hrs)
Untreated
0.1μM DDB/Aplidine
0.5μM DDB/Aplidine
1μM DDB/Aplidine
****
C
293T-ASC-GFP-NLRP1
ASC speck %
ns
***
***
Vector
Vector + ANS
FLAG-SidI WT
FLAG-SidI R453P
B
N/TERT
IL1β (pg/mL)
****
Untreated
0.1 μM
0.5 μM
1 μM
VbP
DDB/Aplidine
D
293T-ASC-GFP-NLRP1
Vector
3xFLAG-SidI
3xFLAG-SidI R453P
FLAG
GAPDH
150
100
75
50
20
37
1
2
3

### Figure S3

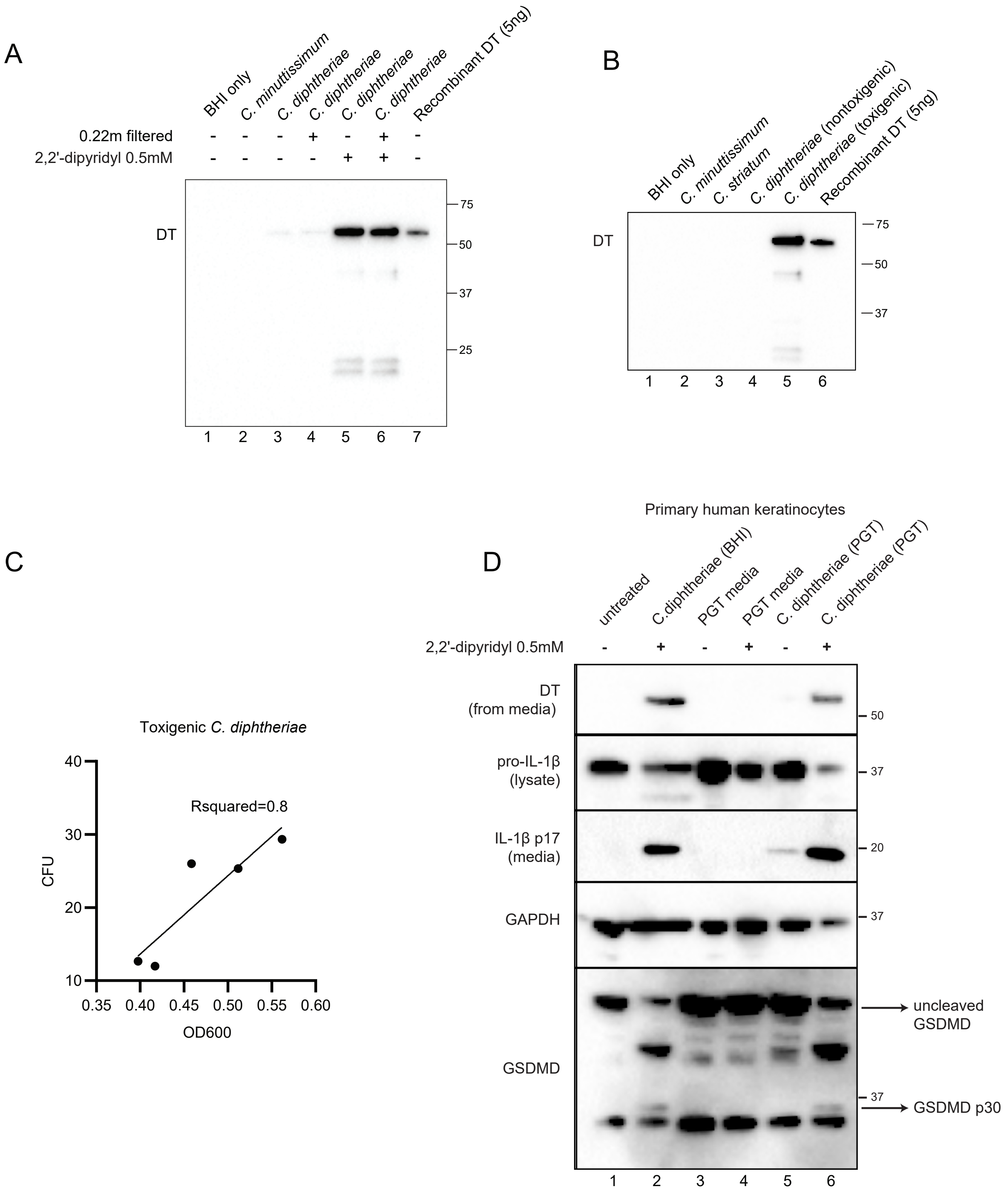
A
BHI only
C. minuttissimum
C. diphtheriae
C. diphtheriae
C. diphtheriae
C. diphtheriae
Recombinant DT (5ng)
0.22m filtered
- - - + - + -
2,2'-dipyridyl 0.5mM
- - - - + + -
DT
75
50
37
25
1 2 3 4 5 6 7
B
BHI only
C. minuttissimum
C. striatum
C. diphtheriae (nontoxigenic)
C. diphtheriae (toxigenic)
Recombinant DT (5ng)
DT
75
50
37
1 2 3 4 5 6
C
Toxigenic C. diphtheriae
Rsquared=0.8
CFU
OD600
10 20 30 40
0.35 0.40 0.45 0.50 0.55 0.60
D
Primary human keratinocytes
untreated
C.diphtheriae (BHI)
PGT media
PGT media
C. diphtheriae (PGT)
C. diphtheriae (PGT)
2,2'-dipyridyl 0.5mM
- + - + - +
DT
(from media)
50
pro-IL-1β
(lysate)
37
IL-1β p17
(media)
20
GAPDH
37
GSDMD
uncleaved
GSDMD
37
GSDMD p30
1 2 3 4 5 6

### Figure S4

A

DPH1 KO validation

| | ICE | KO-Score | KI-Score | ICE d | R Squared |
|---|---|---|---|---|---|
| DPH1 KO (primer 1) | 100 | 99 | None | 100 | 0.75 |
| DPH1 KO (primer 2) | 95 | 95 | None | 86 | 0.93 |

B

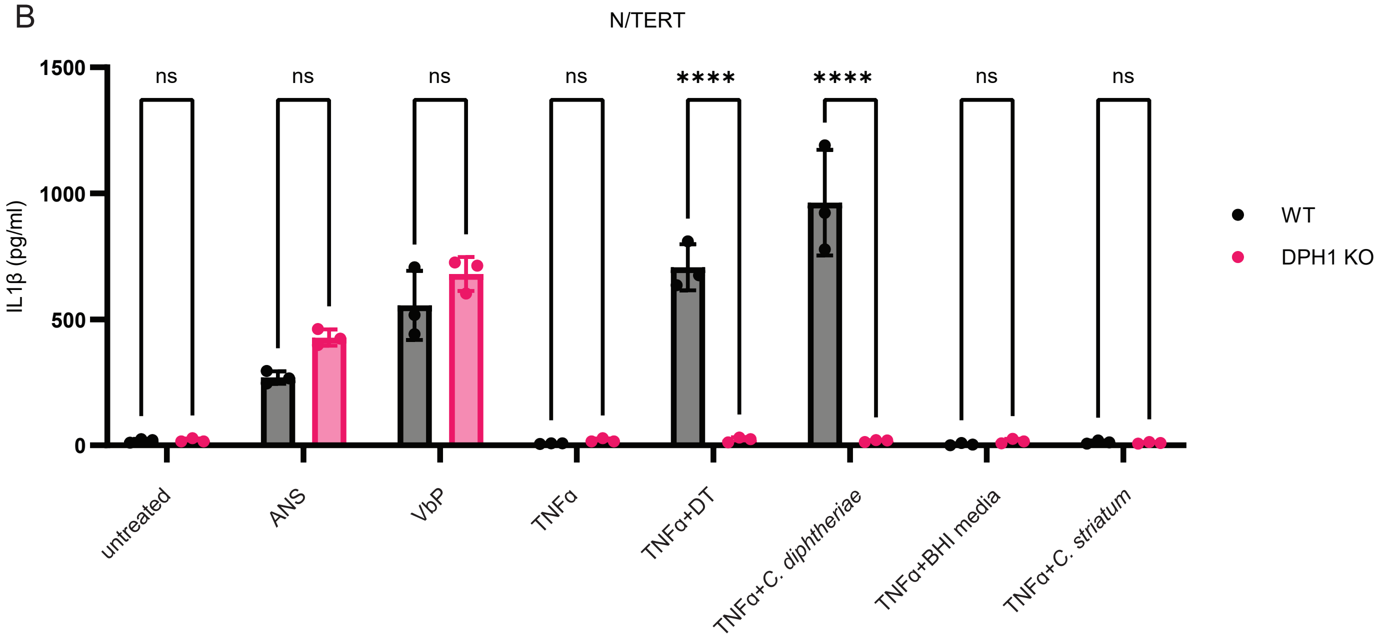

### Figure S5

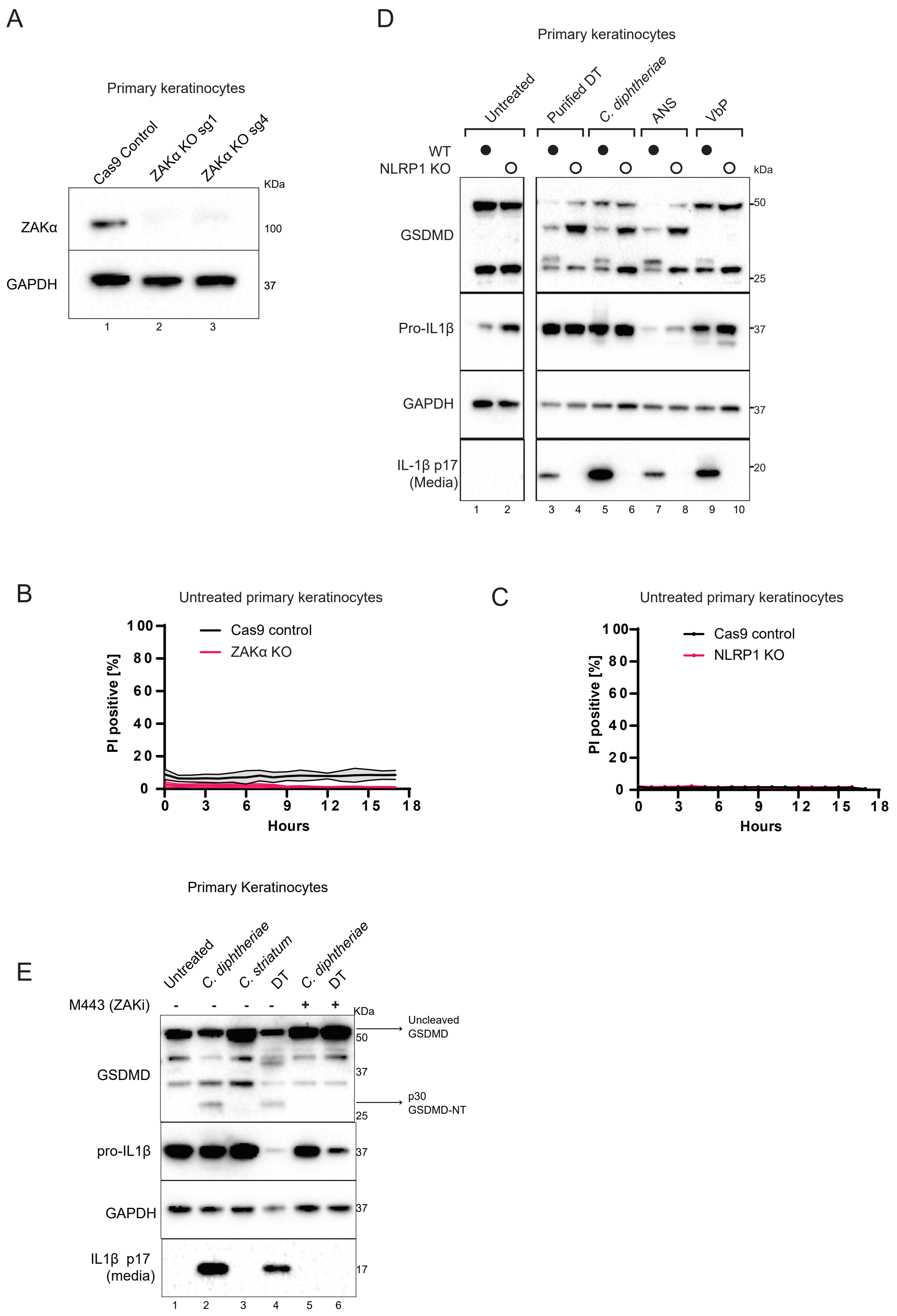
A
Primary keratinocytes
Cas9 Control
ZAKα KO sg1
ZAKα KO sg4
KDa
ZAKα
GAPDH
100
37
1
2
3
D
Primary keratinocytes
Untreated
Purified DT
C. diphtheriae
ANS
VbP
WT
NLRP1 KO
kDa
GSDMD
Pro-IL1β
GAPDH
IL-1β p17
(Media)
50
25
37
37
20
1
2
3
4
5
6
7
8
9
10
B
Untreated primary keratinocytes
Cas9 control
ZAKα KO
PI positive [%]
Hours
0
20
40
60
80
100
3
6
9
12
15
18
C
Untreated primary keratinocytes
Cas9 control
NLRP1 KO
PI positive [%]
Hours
E
Primary Keratinocytes
Untreated
C. diphtheriae
C. striatum
DT
C. diphtheriae
DT
M443 (ZAKi)
KDa
Uncleaved
GSDMD
p30
GSDMD-NT
GSDMD
pro-IL1β
GAPDH
IL1β p17
(media)
50
37
25
37
37
17
1
2
3
4
5
6

### Figure S6

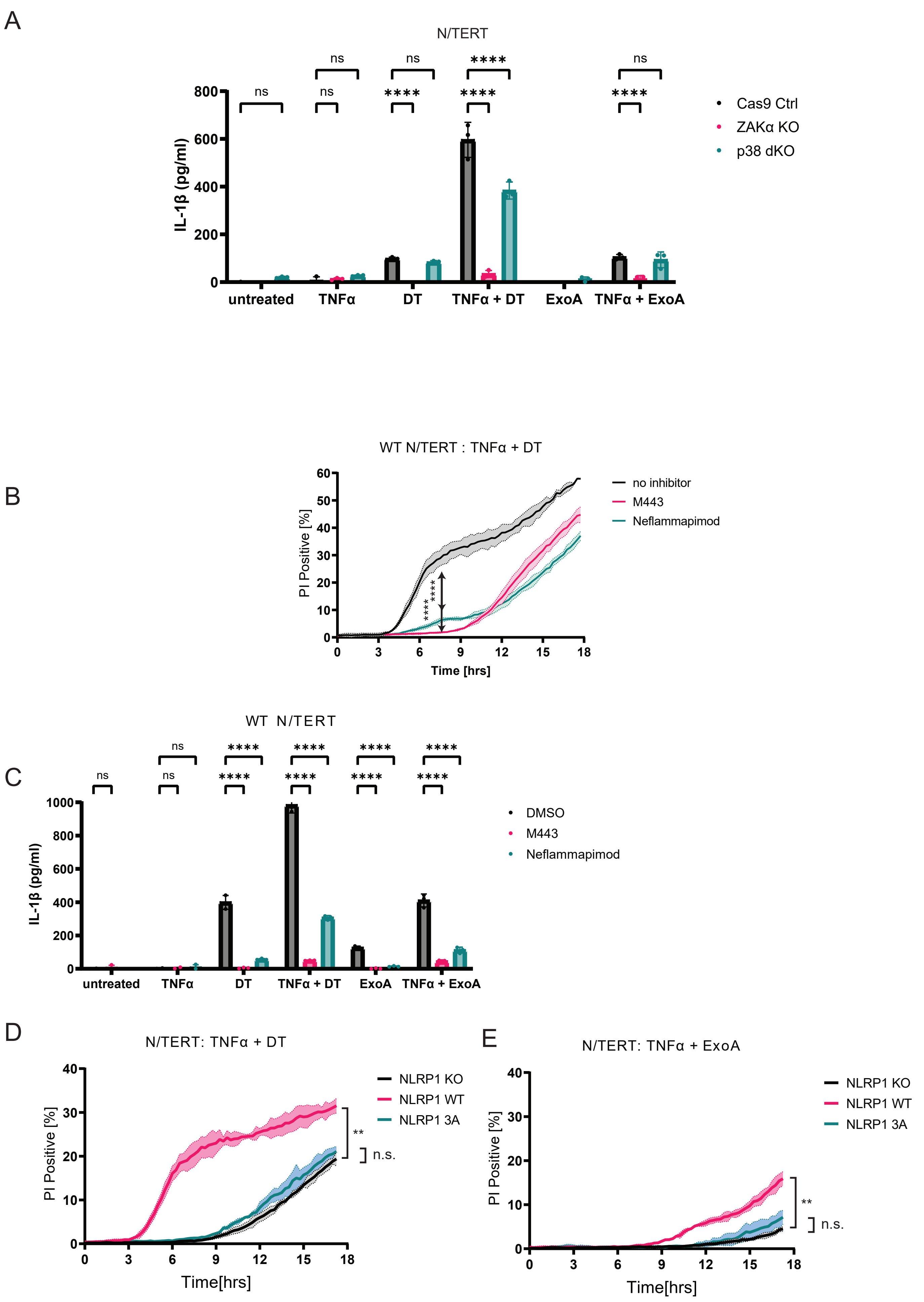

A
N/TERT
IL-1β (pg/ml)
0
200
400
600
800
ns
ns
ns
ns
****
ns
****
****
****
ns
untreated
TNFα
DT
TNFα + DT
ExoA
TNFα + ExoA
Cas9 Ctrl
ZAKα KO
p38 dKO
B
WT N/TERT : TNFα + DT
PI Positive [%]
Time [hrs]
0
3
6
9
12
15
18
0
10
20
30
40
50
60
****
****
no inhibitor
M443
Neflammapimod
C
WT N/TERT
IL-1β (pg/ml)
0
200
400
600
800
1000
ns
ns
ns
****
****
****
****
****
****
****
****
untreated
TNFα
DT
TNFα + DT
ExoA
TNFα + ExoA
DMSO
M443
Neflammapimod
D
N/TERT: TNFα + DT
PI Positive [%]
Time[hrs]
0
3
6
9
12
15
18
0
10
20
30
40
**
n.s.
NLRP1 KO
NLRP1 WT
NLRP1 3A
E
N/TERT: TNFα + ExoA
PI Positive [%]
Time[hrs]
0
3
6
9
12
15
18
0
10
20
30
40
**
n.s.
NLRP1 KO
NLRP1 WT
NLRP1 3A

### Figure S7

A

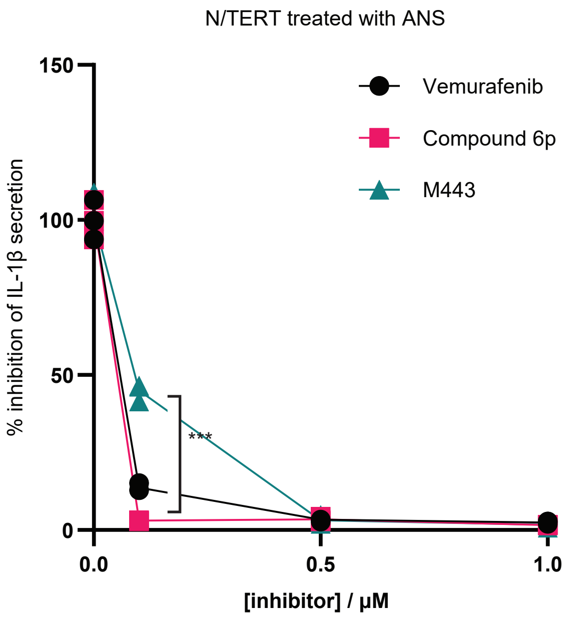


B

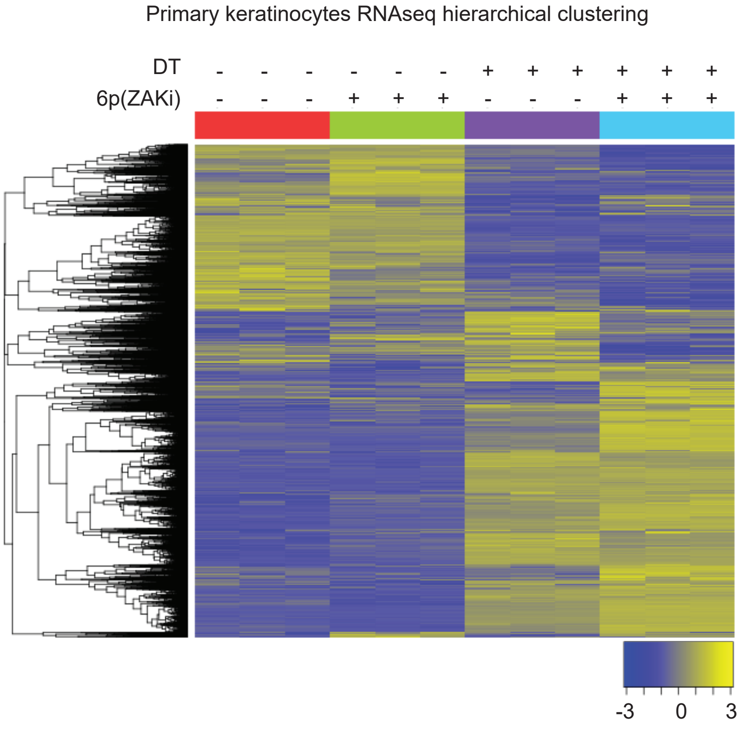


C

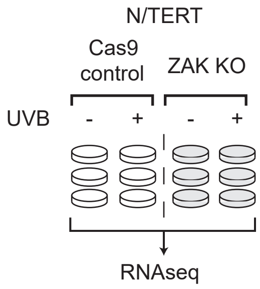


D

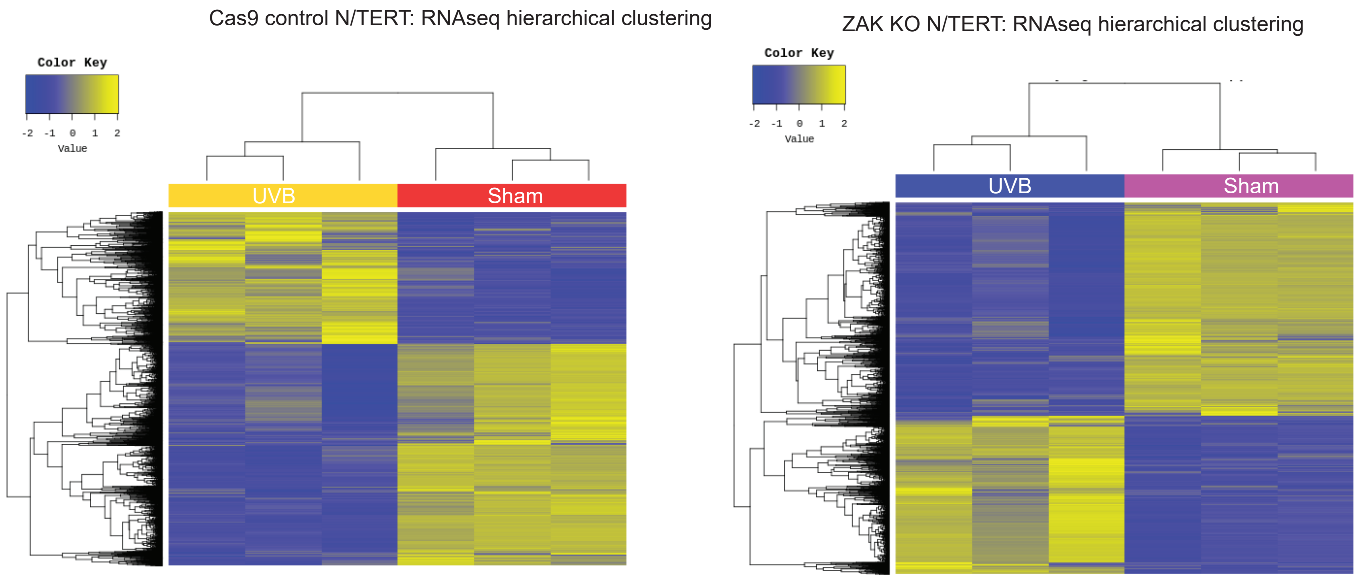

### Figure S8

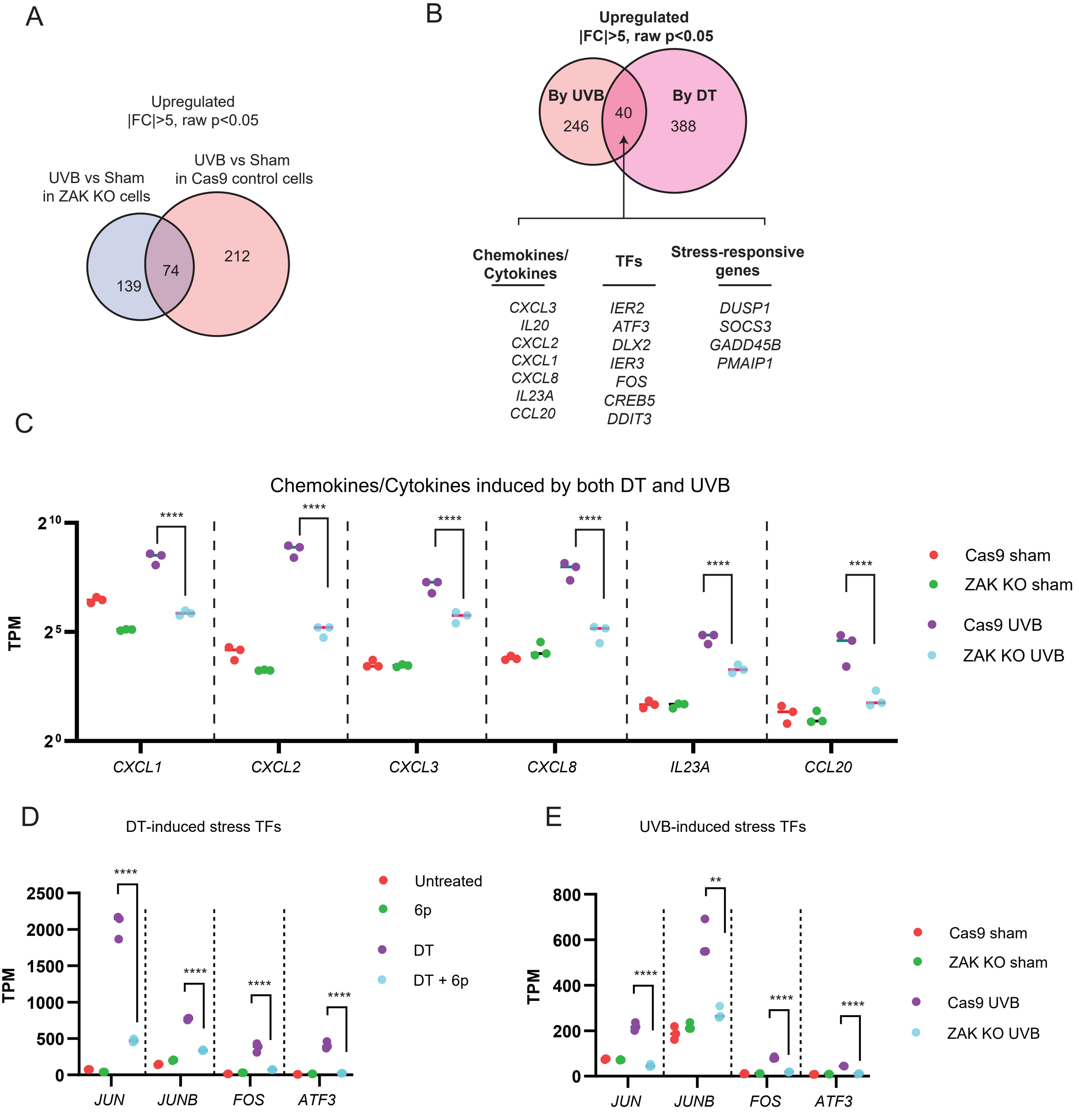
A
Upregulated
|FC|>5, raw p<0.05
UVB vs Sham
in ZAK KO cells
UVB vs Sham
in Cas9 control cells
139
74
212
B
Upregulated
|FC|>5, raw p<0.05
By UVB
By DT
246
40
388
Chemokines/
Cytokines
CXCL3
IL20
CXCL2
CXCL1
CXCL8
IL23A
CCL20
TFs
IER2
ATF3
DLX2
IER3
FOS
CREB5
DDIT3
Stress-responsive
genes
DUSP1
SOCS3
GADD45B
PMAIP1
C
Chemokines/Cytokines induced by both DT and UVB
TPM
$2^{10}$
$2^{5}$
$2^{0}$
CXCL1
CXCL2
CXCL3
CXCL8
IL23A
CCL20
Cas9 sham
ZAK KO sham
Cas9 UVB
ZAK KO UVB
D
DT-induced stress TFs
TPM
2500
2000
1500
1000
500
0
JUN
JUNB
FOS
ATF3
Untreated
6p
DT
DT + 6p
E
UVB-induced stress TFs
TPM
800
600
400
200
0
JUN
JUNB
FOS
ATF3
Cas9 sham
ZAK KO sham
Cas9 UVB
ZAK KO UVB

### Figure S9

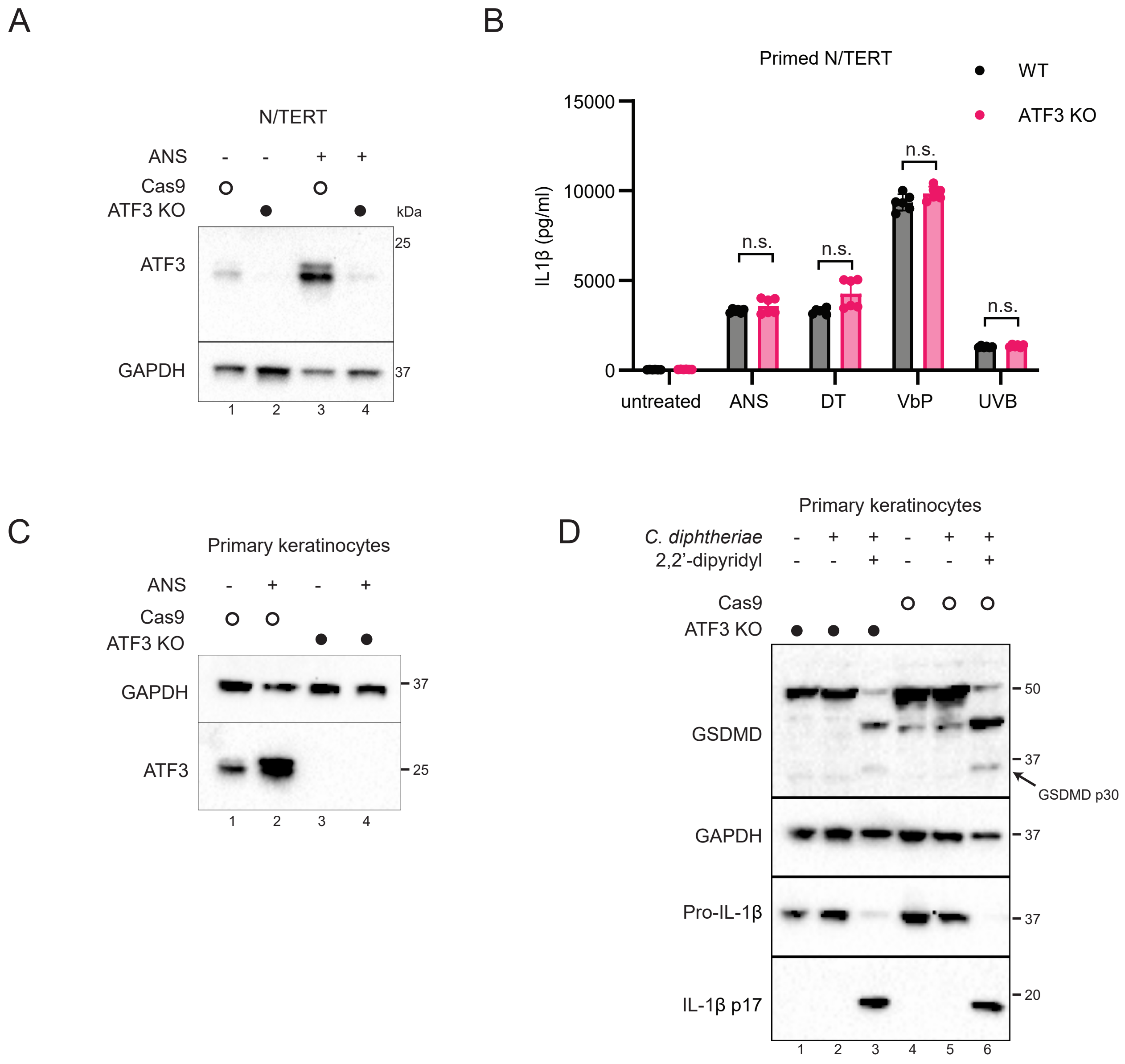
A
N/TERT
ANS - - + +
Cas9
ATF3 KO
kDa
ATF3
25
GAPDH
37
1 2 3 4
B
Primed N/TERT
WT
ATF3 KO
IL1β (pg/ml)
15000
10000
5000
0
n.s.
untreated
ANS
DT
VbP
UVB
C
Primary keratinocytes
ANS - + - +
Cas9
ATF3 KO
GAPDH
37
ATF3
25
1 2 3 4
D
Primary keratinocytes
C. diphtheriae - + + - + +
2,2'-dipyridyl - - + - - +
Cas9
ATF3 KO
GSDMD
50
37
GSDMD p30
GAPDH
37
Pro-IL-1β
37
IL-1β p17
20
1 2 3 4 5 6

### Figure S10

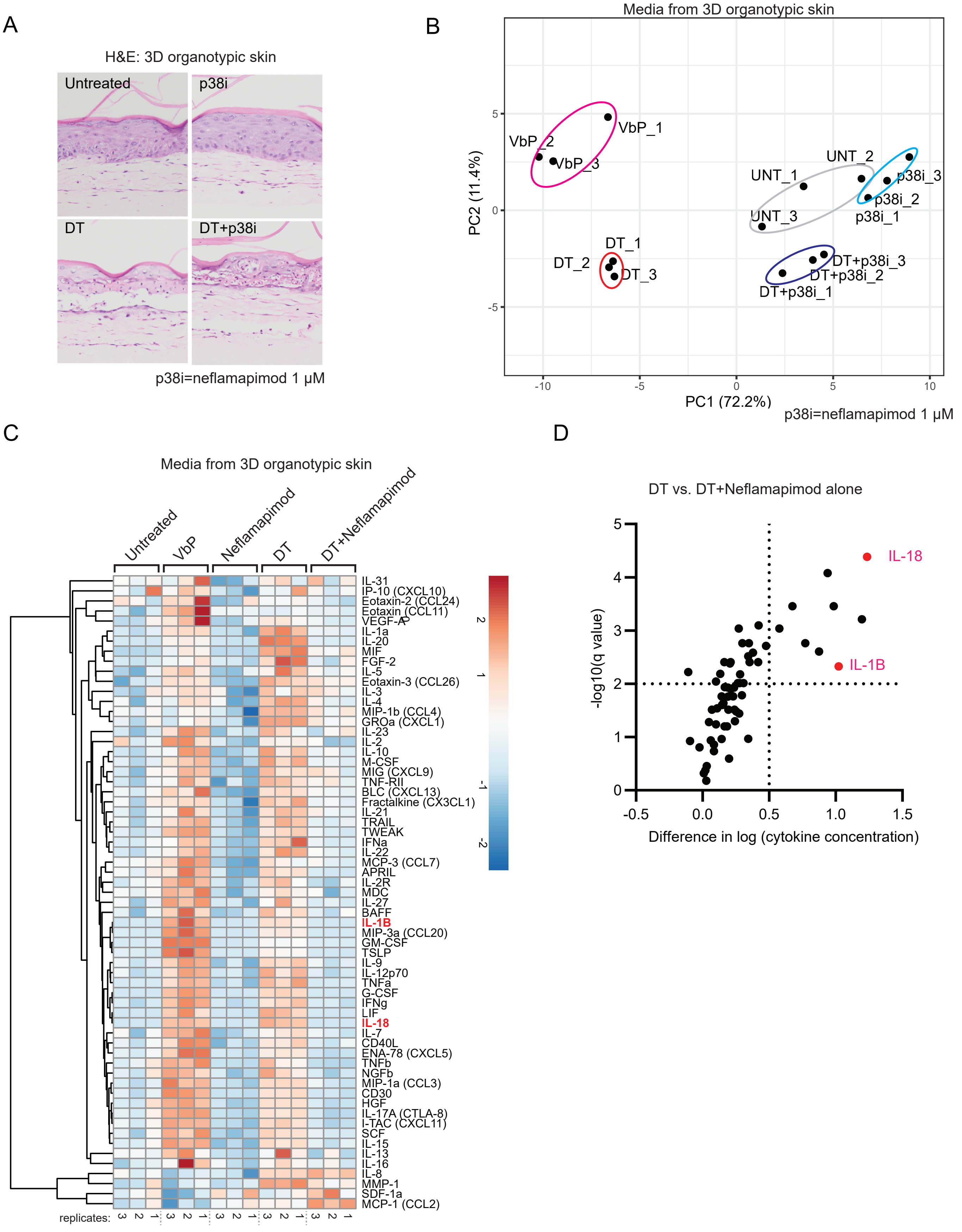
A
H&E: 3D organotypic skin
Untreated
p38i
DT
DT+p38i
p38i=neflamapimod 1 μM
B
Media from 3D organotypic skin
PC2 (11.4%)
PC1 (72.2%)
VbP_1
VbP_2
VbP_3
UNT_1
UNT_2
UNT_3
p38i_1
p38i_2
p38i_3
DT_1
DT_2
DT_3
DT+p38i_1
DT+p38i_2
DT+p38i_3
p38i=neflamapimod 1 μM
C
Media from 3D organotypic skin
Untreated
VbP
Neflamapimod
DT
DT+Neflamapimod
IL-31
IP-10 (CXCL10)
Eotaxin-2 (CCL24)
Eotaxin (CCL11)
VEGF-A
IL-1a
IL-20
MIF
FGF-2
IL-5
Eotaxin-3 (CCL26)
IL-3
IL-4
MIP-1b (CCL4)
GROa (CXCL1)
IL-23
IL-2
IL-10
M-CSF
MIG (CXCL9)
TNF-RII
BLC (CXCL13)
Fractalkine (CX3CL1)
IL-21
TRAIL
TWEAK
IFNa
IL-22
MCP-3 (CCL7)
APRIL
IL-2R
MDC
IL-27
BAFF
IL-1B
MIP-3a (CCL20)
GM-CSF
TSLP
IL-9
IL-12p70
TNFa
G-CSF
IFNg
LIF
IL-18
IL-7
CD40L
ENA-78 (CXCL5)
TNFb
NGFb
MIP-1a (CCL3)
CD30
HGF
IL-17A (CTLA-8)
I-TAC (CXCL11)
SCF
IL-15
IL-13
IL-16
IL-8
MMP-1
SDF-1a
MCP-1 (CCL2)
replicates:
D
DT vs. DT+Neflamapimod alone
-log10(q value)
Difference in log (cytokine concentration)
IL-18
IL-1B

### Figure S11

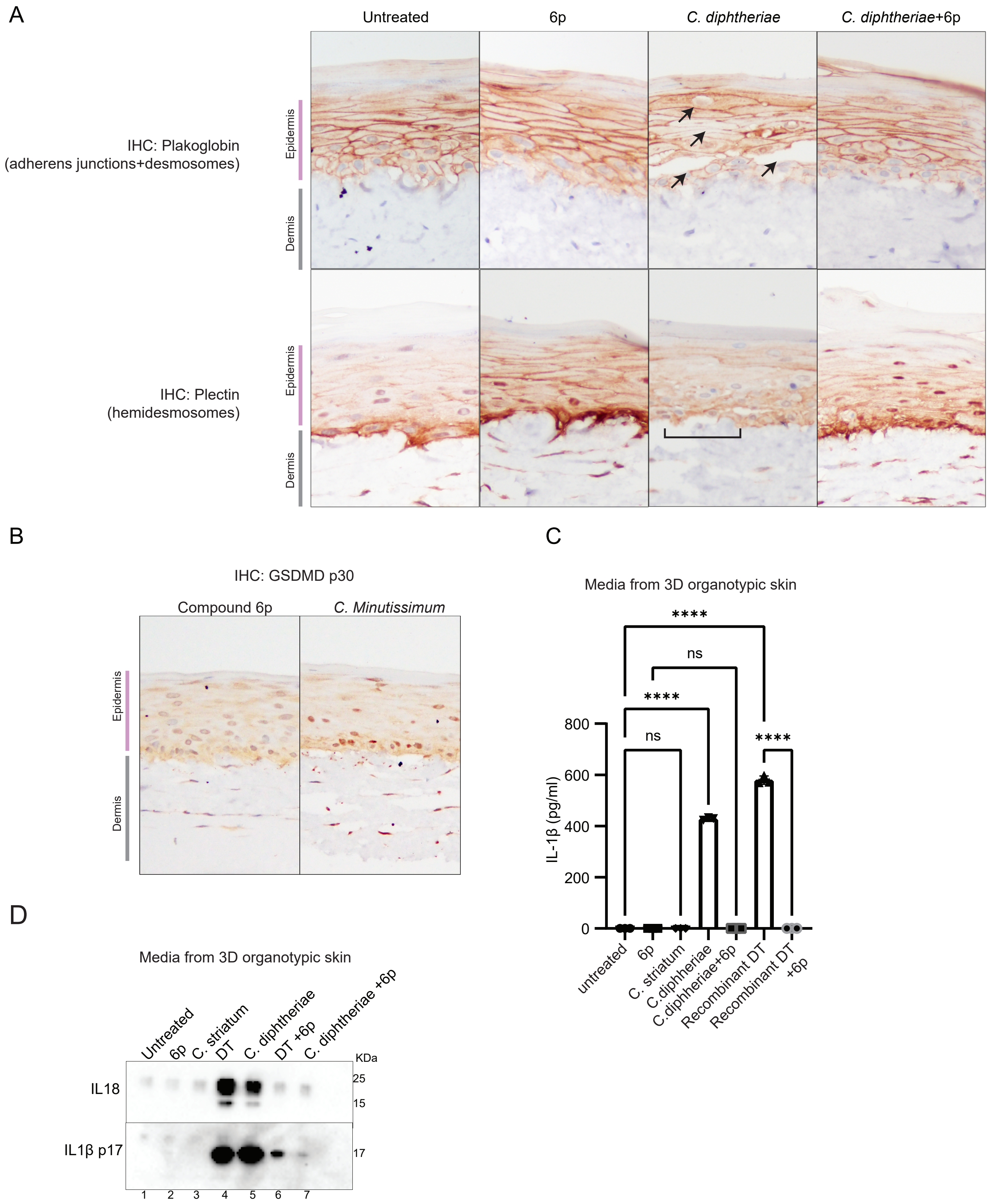

A
Untreated
6p
C. diphtheriae
C. diphtheriae+6p
IHC: Plakoglobin
(adherens junctions+desmosomes)
IHC: Plectin
(hemidesmosomes)
Epidermis
Dermis
B
IHC: GSDMD p30
Compound 6p
C. Minutissimum
Epidermis
Dermis
C
Media from 3D organotypic skin
IL-1β (pg/ml)
0
200
400
600
800
****
ns
****
ns
****
untreated
6p
C. striatum
C.diphtheriae
C.diphtheriae+6p
Recombinant DT
Recombinant DT +6p
D
Media from 3D organotypic skin
Untreated
6p
C. striatum
DT
C. diphtheriae
DT +6p
C. diphtheriae +6p
KDa
IL18
IL1β p17
25
15
17
1
2
3
4
5
6
7
